# An atlas of lamina-associated chromatin across twelve human cell types reveals an intermediate chromatin subtype

**DOI:** 10.1101/2020.07.23.218768

**Authors:** Kathleen C. Keough, Parisha P. Shah, Ketrin Gjoni, Garrett T. Santini, Nadeera M. Wickramasinghe, Carolyn E. Dundes, Ashley Karnay, Angela Chen, Rachel E.A. Salomon, Patrick J. Walsh, Son C. Nguyen, Sean Whalen, Eric F. Joyce, Kyle M. Loh, Nicole Dubois, Katherine S. Pollard, Rajan Jain

**Affiliations:** University of California, San Francisco, CA 94117, USA; Gladstone Institute of Data Science and Biotechnology, San Francisco, CA 94158, USA; Departments of Medicine and Cell and Developmental Biology, Penn CVI, Penn Epigenetics Institute, Perelman School of Medicine, University of Pennsylvania, Philadelphia, PA 19104; Department of Cell, Developmental and Regenerative Biology, Icahn School of Medicine at Mount Sinai, New York, NY 10029; Department of Developmental Biology and Institute for Stem Cell Biology and Regenerative Medicine, Stanford University School of Medicine, Stanford, CA 94305 USA; Department of Genetics, Penn Epigenetics Institute, Perelman School of Medicine, University of Pennsylvania, Philadelphia, PA 19104; Chan Zuckerberg Biohub, San Francisco, CA 94158, USA

**Keywords:** Lamin-associated domains, peripheral chromatin organization, 3D genome, cellular differentiation

## Abstract

Association of chromatin with lamin proteins at the nuclear periphery has emerged as a potential mechanism to coordinate cell type-specific gene expression and maintain cellular identity via gene silencing. Unlike many histone modifications and chromatin-associated proteins, lamin-associated domains (LADs) have yet to be mapped genome-wide in a diverse panel of human cell types, which has limited our understanding of the role peripheral chromatin plays in development and disease. To address this gap, we mapped LAMIN B1 (LB1) across twelve human cell types encompassing pluripotent stem cells, intermediate progenitors, and differentiated cells from all three germ layers. Integrative analyses of this atlas of peripheral chromatin with publicly available genomic data, as well as gene expression and repressive histone maps generated for this study, revealed that in all twelve cellular contexts lamin-associated chromatin is organized into at least two subtypes defined by differences in LB1 occupancy, gene expression, chromatin accessibility, transposable elements, replication timing, and radial positioning. Most genes gain or lose LB1 occupancy consistent with their cell type along developmental trajectories; however, we also identified examples where the enhancer, but not the gene body and promoter, change LAD state. Imaging of fluorescently labeled DNA in single cells validated these transitions and showed intermediate radial positioning of LADs that are gene dense, relatively accessible, and dynamic across development. This atlas represents the largest resource to date for peripheral chromatin organization studies.

## Introduction

Spatial genome organization has emerged as a mechanism for coordinating gene expression towards the specification and maintenance of cellular identity (Rowley and Corces, 2018; Zheng and Xie, 2019). Approximately 30-40% of the genome is localized to the nuclear lamina, a filamentous network of LAMIN A/C, B1, and B2 proteins residing on the inner surface of the nuclear envelope (Burke and Stewart, 2006; Worman and Bonne, 2007). Genomic loci localized at the nuclear lamina, termed lamina-associated domains (LADs), occur in kilobase-to megabase-sized blocks (Guelen et al., 2008; Meuleman et al., 2013). LADs are generally heterochromatic, and genes within LADs are frequently transcriptionally repressed (Briand and Collas, 2020; van Steensel and Belmont, 2017). Mutations in nuclear lamins disrupt chromatin organization and cause disease, and absence of nuclear lamins contributes to chromatin inversion (Briand et al., 2018; Falk et al., 2019; Lee et al., 2019; Shah et al., 2021; Solovei et al., 2013; Vadrot et al., 2015; Worman and Bonne, 2007; Zheng and Xie, 2019). Cellular differentiation has been correlated with repositioning of a subset of genes away from the nuclear lamina and their subsequent expression and vice-versa (Malik et al., 2010; Meister et al., 2010; Peric-Hupkes et al., 2010). More recently, it has been demonstrated that loss of spatial positioning compromises cardiac development (Poleshko et al., 2017). Thus, it is clear that spatial positioning of the genome is important for development and health.

Previous work suggests that a subset of LADs have varying characteristics, such as reduced lamin occupancy and increased gene density, suggesting that LADs are heterogenous across cell types. Comparison of four murine cell types segregated LADs into cell type-variable (facultative) and stable (constitutive) (Kind et al., 2015; Meuleman et al., 2013). Integration of lamin association with core and linker histone occupancy and modifications also distinguished subtypes of LADs within a single murine cell type (Zheng et al., 2015). Subsequent work in single cells showed that the contact frequencies of LADs with the nuclear lamina vary by locus and single cell studies suggest that individual genomic regions, in aggregate, have varying probabilities of becoming re-localized to or from the nuclear lamina during differentiation (Kind et al., 2013, 2015; Rooijers et al., 2019), leading to the model predicting different subtypes of LADs. Our understanding of LAD subtypes, however, is still far from complete due to the limited number of contexts that have been studied to date. Further study of lamina-associated chromatin across a greater diversity of cell types and identification of key molecular characteristics will provide critical knowledge about how LADs are organized and clues into how spatial positioning may contribute to cellular identity.

To address these gaps, we developed an atlas of LAMIN B1 (LB1) binding across twelve human cell types from all three germ layers and embryonic stem cells (ESCs). Our data revealed two subtypes of lamina-associated chromatin in all examined cell types, one of which represents LADs with intermediate LB1 enrichment, chromatin accessibility, dimethylation of lysine 9 on histone H3 (H3K9me2), gene density, and gene expression. Unlike previous approaches, our modeling allowed for identification of different LAD subtypes within individual cell types, without the requirement for comparisons between cell types. Imaging in two cell types demonstrated a difference in radial positioning of the two identified subtypes of LADs. The intermediate LAD subtype is also more dynamic across cell types, with cell type-specific gene expression, differential chromatin accessibility, and differentially active enhancers. Overall, this work provides the largest atlas of human lamina-associated chromatin maps to date, reveals important insights into how association with the lamina may participate in cell type specification and development, and provides an important resource for future interrogations of peripheral chromatin.

## Results

### A three-state hidden Markov model identifies two subtypes of LADs in 12 human cell types

We generated an atlas of LB1 occupancy from human ESCs and twelve ESC-derived cell types from all three germ layers (endoderm, mesoderm and ectoderm) using established and published differentiation protocols representative of early developmental trajectories (Ang et al., 2018; Bardot et al., 2017; Loh et al., 2014, 2016; Martin et al., 2020) (Fig. 1A; Table 1). Expression of key genes by quantitative reverse transcription PCR (qRT-PCR, Supp. Fig. 1A-F), fluorescence-activated cell sorting, and immunofluorescence for cell type-specific factors (Supp. Fig. 1G-M) in various cell types supported our established differentiation strategies (Ang et al., 2018; Dubois et al., 2011; Josowitz et al., 2014; Loh et al., 2014, 2016). Cardiac cultures also displayed stereotypical contractile activity (Supp. Movie 1, 2). We performed LB1 chromatin immunoprecipitation (ChIP) using an antibody that has been previously validated for ChIP in human cells (Sadaie et al., 2013; Shah et al., 2013, 2021). We further verified the antibody in select cell types by performing immunoblot following LB1 ChIP under standard conditions and with the addition of a LB1 peptide, which quenched LB1 immunoprecipitation (Supp. Fig. 2A, B). We chose ChIP-seq, rather than DamID, to map LB1 across multiple stages of differentiation, because it reflects targeted protein binding at fixation, versus DNA methylation that occurs over several hours and circumvents differences in methylation efficiency or differentiation that may occur when using DAM-modified cells (Aughey et al., 2019; Rooijers et al., 2019; van Steensel and Henikoff, 2000). Each biological replicate was sequenced to greater than 40 million uniquely mapping reads, and the Pearson correlation between replicates was greater than 0.7 for 90% of cell types, indicating high reproducibility between replicates (Table 1). Visual inspection of the atlas LB1 ChIP-seq data on a genome browser confirmed the presence of large, discrete LB1-enriched domains consistent with the presence of LADs in all cell types investigated (Fig. 1B).

**Fig. 1:**
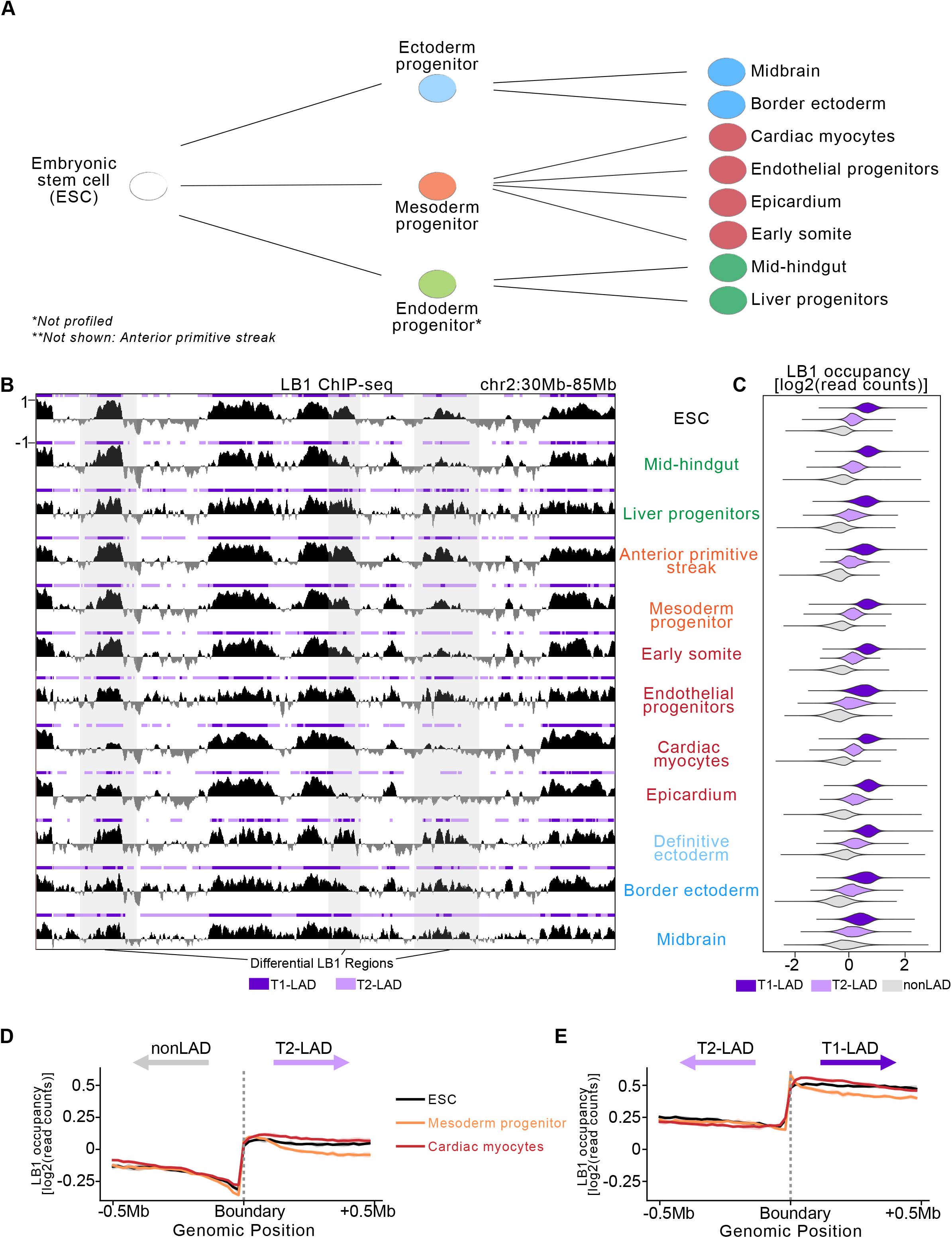
HMM identifies 3 state LADs from LB1 ChIPseq datasets across 13 human cell types. A. Simplified overview of progressive lineage restriction indicating some of the cell types represented in the LAD atlas, shown from greatest (left) to least (right) developmental potential. B. Example track view of LB1 enrichment and associated LADs in all 12 cell types assayed (representative replicate track shown; Y-axis scale is the same in all datasets). LAD states defined by HMM per cell type shown above each LB1 track. T1-LADs in dark purple; T2-LADs in light purple. Regions highlighted in grey span differential LB1/LAD regions between cell types. Y-axis indicated on the first track and consistent across all tracks shown. C. LB1 occupancy per LAD state in all 12 cell types. T1-LADs (dark purple) are the most enriched and nonLADs (grey) are least enriched for LB1. D,E. LB1 occupancy measured 500kb up and downstream of boundaries between (D) nonLADs and T2-LADs (including mirrored T2-LAD to nonLAD boundaries) and (E) T2-LADs and T1-LADs (including mirrored T1-LAD to T2-LAD boundaries) shows sharp distinction of LB1 signal between these features.

We implemented hidden Markov models (HMMs) to identify LADs in each cell type using the LB1 ChIP-seq datasets as input (see Methods). We chose HMMs over broad peak callers like Enriched Domain Detector (Lund et al., 2014) to circumvent the necessary manual parameter tuning across such a large dataset. In addition, such broad peak callers often only identify the LAD and nonLAD state. HMMs have previously been used to segment chromosomes into LAD versus nonLAD regions utilizing DamID-generated LB1 binding peaks, and to segment chromatin states based on various epigenomic marks (Ernst and Kellis, 2017; Filion et al., 2010; Meuleman et al., 2013). We compared model fits for two through five states to determine how many distinct types of domains were contained in the data (Supp. Fig. 3A, B). Our goal was to fit the simplest model that accurately describes the data. The improvements in model fit with three as opposed to two states suggested the existence of an intermediate state of lamina-associated chromatin in all 12 human cell types, consistent with previous observations of an intermediate LAD state in a single cell type studied (Zheng et al., 2015). We therefore proceeded with a 3-state HMM for each cell type.

We designated the LB1-defined states as Type 1 LADs (T1-LADs), Type 2 LADs (T2-LADs) and nonLADs in descending order of LB1 occupancy (Fig 1B, C, Table 2). T1-LADs are slightly larger [median across cell types: 290 kilobases (kb)] than T2-LADs (median: 280kb), though domain sizes vary greatly for both subtypes [T1 range: 20kb-10.5 megabases (Mb), T1 IQR: 620kb, T2 range: 20kb-9.3Mb, T2 IQR: 440kb]. T1-LADs and T2-LADs respectively cover 31.5% and 35.3% of the genome on average across cell types. We assessed LB1 occupancy across 20 different T1-LAD, T2-LAD, and nonLAD regions by LB1 ChIP-qPCR which verified differential LB1 enrichment (Supp. Fig. 3C). It has been previously shown (Shah et al, 2013; Poleshko et al, 2017) that ChIP-seq-generated LADs have a high degree of overlap with DamID-generated LADs. As expected, we found greater than 90% of DamID LB1-positive probes from human ESCs (Meuleman et al., 2013) represented in T1- or T2-LADs from ESCs profiled in this study (Supp. Fig. 3D). Moreover, LB1 occupancy across nonLAD:T2-LAD (Fig. 1D) and T1-LAD:T2-LAD (Fig. 1E) transitions showed sharp changes in LB1 enrichment with different levels of LB1 binding on either side of the boundary in select cell types that were used for further interrogation (see below), indicating distinct transitions between the LAD states. Thus, using LB1 ChIP-seq and cell type-specific 3-state HMMs, we identified two subtypes of LADs in 12 human cell types.

### T1- and T2-LADs have distinct genomic features

Given their differential LB1 enrichment, we hypothesized that T1- and T2-LADs would differ in other epigenomic features. We first assessed LADs for occupancy of H3K9me2, a histone modification enriched in lamina-associated chromatin (Kind et al., 2013; Poleshko et al., 2017; See et al., 2019; Shah et al., 2021). We generated H3K9me2 ChIP-seq maps (Fig. 2A) using an antibody we previously extensively validated for specificity for H3K9me2 with modified histone peptide arrays (Poleshko et al., 2017; Shah et al., 2021). H3K9me2 ChIP-immunoblot (Supp. Fig. 4A) and peptide immunoblot (Supp. Fig. 4B) also indicated specificity for H3K9me2 as opposed to closely related modifications. We applied similar HMM fitting procedures to the H3K9me2 data, again finding a 3-state model to be the simplest model that captures a large proportion of the variability in our ChIP-seq data (Fig. 2A; Supp. Fig. 5A, B). These states were designated T1-, T2-, and nonKDDs (<u>K</u>9 <u>d</u>imethyl <u>d</u>omains) based on decreasing H3K9me2 occupancy (Supp. Fig. 5C, Table 3), which we validated by H3K9me2 ChIP-qPCR (Supp. Fig. 5D). Across cell types, T1- and T2-KDDs demonstrated a high degree of overlap with T1- and T2-LADs, respectively (Supp. Fig. 5E, F). Accordingly, LB1 occupancy was most often highest in T1-KDDs, followed by T2-KDDs and nonKDDs (Fig. 2B). This analysis confirmed that in 12 human cell types, greater chromatin association with LB1 coincides with increasing H3K9me2 occupancy.

**Fig. 2:**
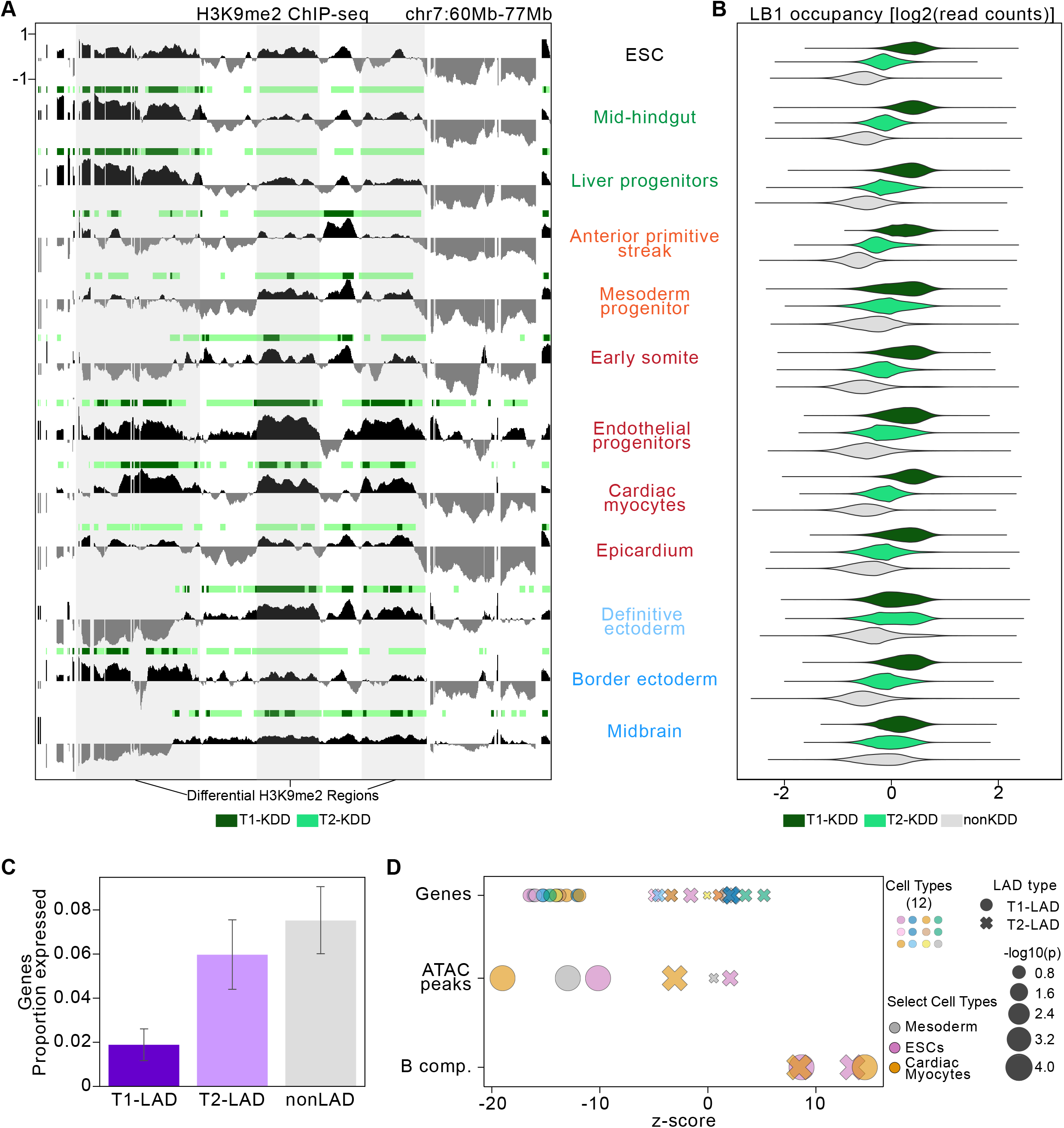
LADs correspond closely with KDDs and demonstrate characteristic features across cell types. A. Example track view of H3K9me2 enrichment and associated KDDs in all 12 cell types assayed. KDD states as defined by HMM per cell type shown above each H3K9me2 track. T1-KDDs in dark green; T2-KDDs in light green. Regions highlighted in grey span differential H3K9me2/KDD regions between cell types. Y-axis indicated on the first track and consistent across all tracks shown. C. LB1 occupancy per KDD state in all 12 cell types. T1-KDDs (dark green) are the most enriched and nonKDDs (grey) are least enriched for LB1. C. Mean proportion of genes expressed (TPM > 5) per LAD category for cell types with matched RNA-seq data (cardiac myocytes, early somite, mesoderm progenitors, mid-hindgut, definitive ectoderm and endothelial progenitors). Error bars indicate standard deviation. D. Enrichment of genes, ATAC-seq peaks (Liu et al. 2017), and B compartment (B comp.) from matched cell types (Zhang et al. 2019, see Methods). Cell types in key are as follows: Row 1 (L-R): ESC, liver progenitors, epicardium, endothelial progenitors; Row 2 (L-R): mid-hindgut, border ectoderm, anterior primitive streak, midbrain; Row 3 (L-R): cardiac myocytes, definitive ectoderm, early somite, mesoderm progenitors.

We next investigated additional characteristic features of lamin-associated chromatin across cell types. Compared to T2-LADs and nonLADs, T1-LADs had the lowest proportion of expressed genes based on RNA-seq generated for this study and from published sources from matched cell types (Fig. 2C) (Koh et al., 2016; Loh et al., 2014; Tchieu et al., 2017). While both types of LADs are heterochromatic, as evidenced by enrichment for B compartment regions (Lieberman-Aiden et al., 2009) (Fig. 2D), T1-LADs showed the greatest depletion for genes and ATAC-seq peaks (Liu et al., 2017), while T2-LADs were consistently intermediate between T1-LADs and nonLADs for gene expression, gene density, and accessibility across multiple cell types (Fig. 2C-D). Motivated by these results, we sought to better understand distinguishing characteristics between T1- and T2-LADs. We first determined that specific types of transposable elements show divergent patterns of enrichment in T1- versus T2-LADs (Fig. 3A, upper), though L1 and ERVL-MaLR elements are consistently enriched in both LAD subtypes. Next, though T1- and T2-LADs were depleted for constitutively early replicating domains compared to nonLADs (Dixon et al., 2018), we found that only T1-LADs were enriched for constitutive late replication timing domains, while T2-LADs were enriched for switch replication timing domains (Fig. 3A, lower). In addition, we assessed binding sites of the chromatin insulator CTCF from comparable cell types (Zhang et al., 2019) and observed that CTCF occupancy decreased sharply at transitions from states of lower to higher LB1 binding (from nonLAD to T2-LAD or T2-LAD to T1-LAD) following an increase at the boundary itself (Fig. 3B, C), consistent with previous results (Guelen et al., 2008; Handoko et al., 2011; Harr et al., 2015; van Schaik et al.). T1-LADs also had the most overlap with all LB1-associated regions defined by single cell DamID (Kind et al., 2013) in a comparable atlas cell type, suggesting that at least a subset of T2-LADs may be more variable in a population of cells (Fig. 3D).

**Fig. 3:**
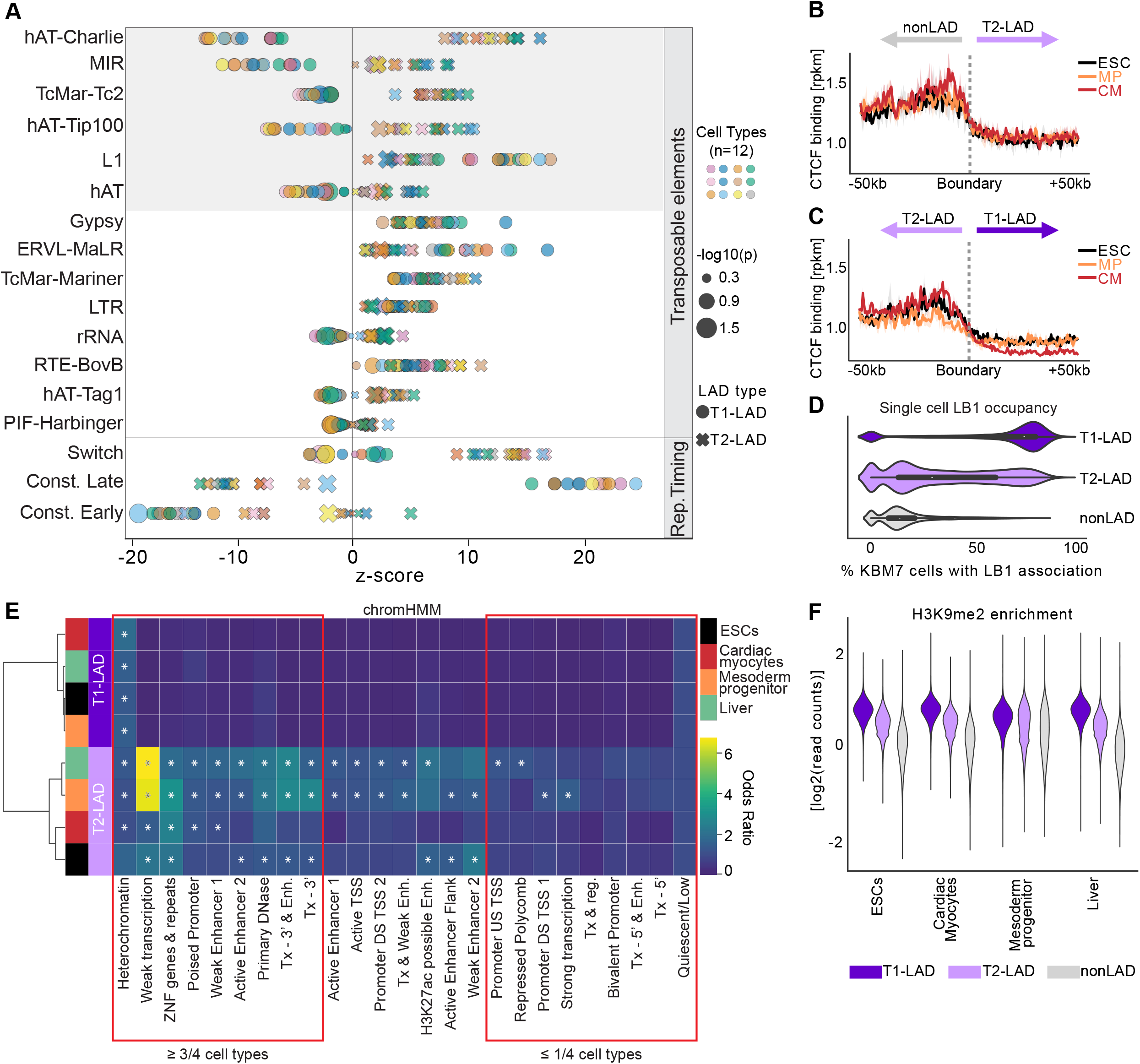
Genomic characterization of LAD subtypes indicates T1- and T2-LAD distinction. A. Enrichments of transposable elements (top) and replication timing (RT) domains (bottom) in T1- and T2-LADs across the atlas. Transposable elements differentially enriched in T1- and T2- LADs highlighted in grey. Cell types in key are as follows: Row 1 (L-R): ESC, liver progenitors, epicardium, endothelial progenitors; Row 2 (L-R): mid-hindgut, border ectoderm, anterior primitive streak, midbrain; Row 3 (L-R): cardiac myocytes, definitive ectoderm, early somite, mesoderm progenitors. B,C. CTCF binding at transitions from (B) nonLADs to T2-LADs (including mirrored T2-LAD to nonLAD boundaries) and (C) T2-LADs to T1-LADs (including mirrored T1-LAD to T2- LAD boundaries) shows CTCF enrichment at T2-LAD boundaries. D. Percentage of single cells with LB1 occupancy per LAD locus in mesoderm progenitor cells shows greater overlap with T1- LADs. E. chromHMM feature enrichments in LADs from four cell types, as indicated. Asterisks (*) indicate adjusted p-value < 0.01. F. H3K9me2 occupancy per LAD state in selected cell types, as indicated (full set in Supp. Fig. 5G). T1-LADs (dark purple) are the most enriched and nonLADs (grey) are least enriched for H3K9me2.

In order to integrate the findings, we turned to chromHMM, a platform that can annotate different states based on various epigenetic and related characteristics (Ernst and Kellis, 2017). T1- and T2-LADs were enriched for the chromHMM heterochromatin state (Ernst and Kellis, 2017) in all cell types, except for T2-LADs in ESCs, which did not meet statistical significance. Overall, T2-LADs harbored a more diverse group of states than T1-LADs. Most notably, T2-LADs were consistently depleted of bivalent promoters, repressed polycomb chromatin, strong transcription regions, and related states (at least 3 of 4 cell types). Conversely, T2-LADs were significantly enriched in all cell types assessed for weak transcription, and zinc finger genes and repeats in all four cell types and significantly enriched for multiple enhancer states in at least three of four cell types (Fig. 3E). Consistent with the chromHMM output and the close alignment of LB1 and H3K9me2, we found that most often H3K9me2 showed the highest enrichment in T1-LADs, followed by T2-LADs and then nonLADs (Fig. 3F, Supp. Fig. 5G). Collectively, these findings of divergent enrichment patterns for epigenetic states, transposable elements and replication timing domains, strong boundary signals from CTCF, and differing patterns in single cell DamID data suggest that T1- and T2-LADs are distinct entities.

### T2-LADs are distinctly spatially positioned from T1-LADs

The above analyses raised the possibility that T1- and T2-LADs may be differentially localized within the nucleus. Our data predicted that T1-LADs would be spatially positioned at the nuclear lamina, given that this LAD subtype has the highest LB1 occupancy. It was unclear, however, whether any given T2-LAD would be heterogeneously positioned with respect to the periphery across cells, as suggested by single cell Dam-ID and related studies (Kind et al., 2015; Rooijers et al., 2019). Alternatively, it was plausible that an individual T2-LAD would be consistently positioned farther away from the nuclear lamina compared to T1-LADs, similar to a recent integrated analysis of Dam-ID and TSA-seq data from K562 cells, which suggested that loci consistently fall into specific locations within the nucleus (Wang et al., 2021). To test these predictions and distinguish between these possibilities, we performed immunofluorescence of LB1 coupled with oligo-based fluorescence *in situ* hybridization (IF-FISH) (Beliveau et al., 2012) to visualize the nuclear lamina and multiple T1-LAD, T2-LAD, and nonLAD regions (Fig. 4).

**Fig. 4:**
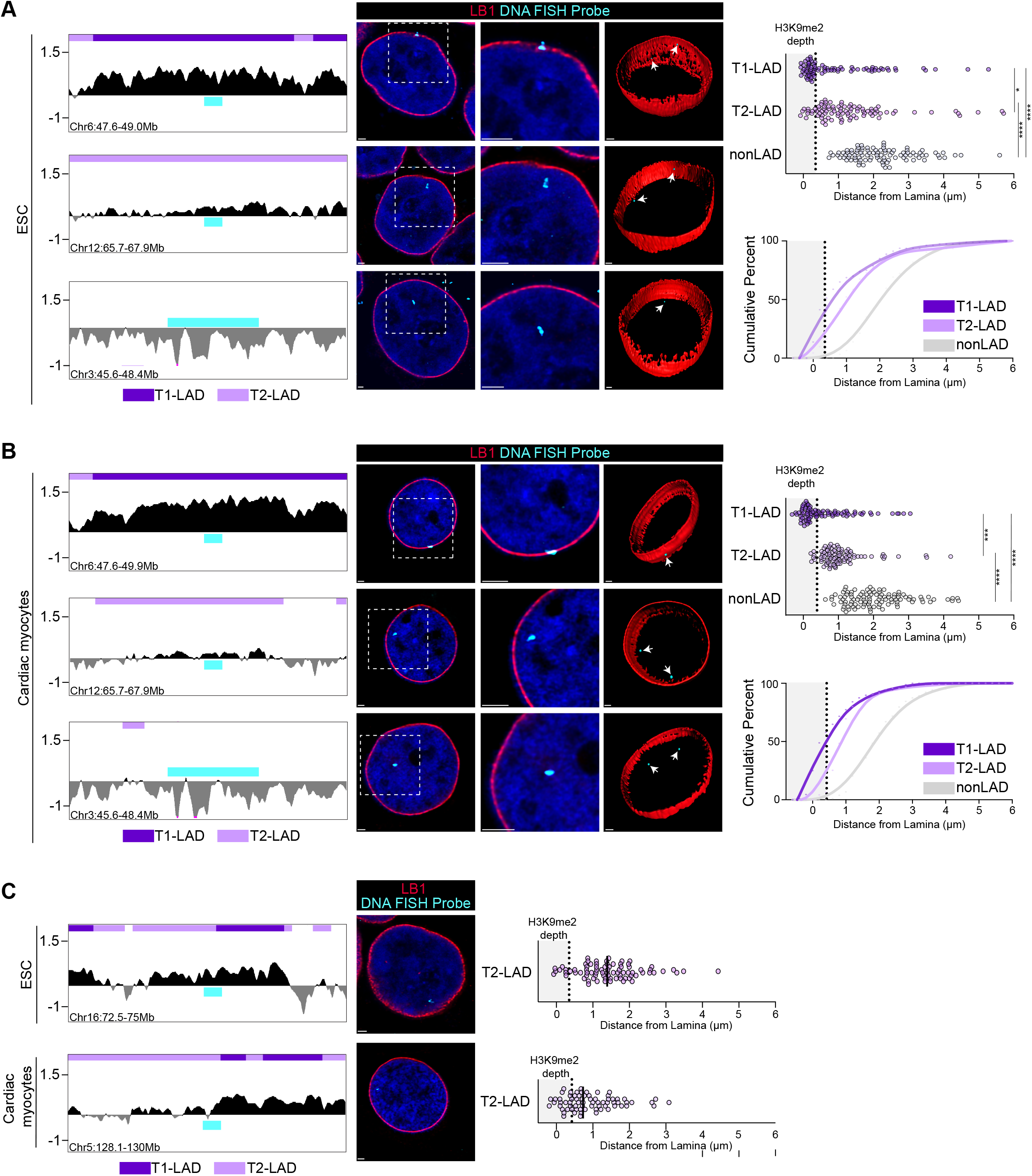
T1-LAD, T2-LAD and nonLAD regions are distinctly radially positioned within the nucleus. A,B. (Left) T1-LAD, T2-LAD, and nonLAD FISH probe locations in LB1 tracks in ESCs (A) and cardiac myocytes (B). (Middle) Representative images of FISH foci (individual slices from Z-stack) from each tested probe region with 3D renderings of all slices to the right for ESCs (A) and fluorescence activated cell sorted cardiac myocytes (B). Box in the left image indicates the approximate area of the high magnification image in the middle. (Right) Quantification (see Methods) of the distance of each FISH focus per probe relative to the middle of the nuclear lamina (LB1) signal shown by total (top) and cumulative frequency distribution (bottom). Grey boxes in the quantification plots indicate the 90th percentile depth of H3K9me2-marked heterochromatin at the nuclear periphery (see Methods and Supp. Fig. 6A). Statistical significance of the differences in localization between T1-, T2-, and nonLAD foci calculated by a Kruskal-Wallis test with post-hoc Dunn test for multiple comparisons. *p<0.05, ***p<0.001, ****p<0.0001. C. Probe tracks (left), representative images (middle) and quantification (right) of additional probes primarily targeting T2-LADs in ESCs (top) and in cardiac myocytes (bottom). Black bar in quantification plots marks the median. Mann Whitney U test was performed to compare each additional probe set in C to the cell-type and LAD type matched probe in A or B; no significant differences were observed for both comparisons. Scale bars = 1µM.

We first selected three candidate loci classified as T1-LAD, T2-LAD and nonLAD in both ESCs (Fig. 4A) and cardiac myocytes (Fig. 4B). We performed IF-FISH and measured the shortest 3D Euclidean distance of the FISH foci to the nuclear lamina (see Methods). As predicted, the T1- LAD foci were often embedded within the nuclear lamina and the nonLAD foci were largely excluded from the nuclear lamina (Fig. 4A, right; n=46 and 49 nuclei for T1- and nonLAD, respectively). The majority of foci for the T2-LAD probed region, however, were positioned just adjacent to the nuclear lamina (Fig. 4A, right; n=49 nuclei). We also confirmed the enrichment of H3K9me2-marked chromatin at the nuclear periphery in ESCs by IF as a proxy for the chromatin localized at the nuclear periphery (Supp. Fig. 6A). The depth of H3K9me2-marked chromatin was consistent with previous studies in murine cells (Poleshko et al., 2017). We found 51% of T1-LAD foci within the 90th percentile of H3K9me2 depth (90th percentile depth = 349nm based on 5 independent measurements of the inner/outer LB1 signal to edge of H3K9me2 signal in n=10 independent nuclei - see Methods; Supp. Fig. 6A), compared to only 16% T2-LAD foci. As expected, no nonLAD foci fell within this threshold (Fig. 4A, right).

We next performed IF-FISH with the same probes in cardiac myocytes. For these experiments, we utilized fluorescence-activated cell sorting to generate pure populations of cardiac myocytes (Supp. Fig. 6B). Strikingly, the IF-FISH results were nearly identical between cardiac myocytes and ESCs (Fig. 4B; n=51, 49, and 55 nuclei for T1-, T2-, and nonLAD, respectively), with the majority of T2-LAD probed regions specifically positioned just adjacent to the nuclear lamina. After measuring the depth of the H3K9me2-marked chromatin as above (90% percentile depth = 422nm; Supp. Fig. 6A), we observed that 58% of T1-LAD foci were within this threshold, compared to only 6% of T2-LAD foci, with most T2-LAD foci localized just adjacent to the nuclear lamina. IF-FISH to an additional T1-LAD locus in cardiac myocytes also localized to the nuclear lamina, with most foci in the H3K9me2-marked chromatin (Supp. Fig. 6C, n=43 nuclei). No nonLAD foci were localized within this threshold (Fig. 4B, right), as in ESCs.

To expand these results, we performed IF-FISH against additional loci which were predominantly T2-LADs in ESCs (n=49 nuclei) and cardiac myocytes (n=42 nuclei; Fig. 4C); quantification of FISH foci localization supported the general distinction between LAD and nonLAD localization. Notably, each probe is mostly within a T2-LAD with partial overlap in a T1-LAD; concordantly, we observed an overall distribution of foci within the T2-LAD range, but with a greater portion of foci within the 90% percentile threshold for each compared to the fully T2-LAD probe above (Fig. 4A, B). These data suggested that the reduced relative LB1 occupancy as measured by ChIP-seq in the multiple candidate T2-LAD regions is more likely explained by radial positioning in weak LB1 chromatin regions at the nuclear periphery as opposed to being the result of dynamic positioning across a population of cells. The IF-FISH data, taken collectively with the molecular characteristic assessments above, indicate that T1-LAD and T2-LADs are distinct features that can be defined within and across cell types and that have distinct radial positioning, with T2-LADs occupying an intermediate LAD space between T1-LADs and nonLADs.

### Most T1- and T2-LADs vary across cell types

Previous studies have shown that many LADs are shared between cell types (Meuleman et al., 2013; Robson et al., 2016); however, these studies are often limited to comparisons between two or three cell types along a single differentiation trajectory. We therefore hypothesized that invariant LADs, genomic regions which are conserved as the same LAD subtype in all the atlas datasets, would be the exception rather than the rule in our atlas of 12 cell types. Indeed, we identified only a small fraction of the genome as invariant T1- and T2-LADs, 7% and 1% of the human genome, respectively, with representative chromosomes shown in Fig. 5A. An example of a gene located in an invariant T1-LAD is *OR10K1*, an olfactory receptor gene (Fig 5B). Odorant receptor genes are highly repressed except in olfactory sensory neurons, where multiple enhancers facilitate the expression of a single odorant receptor allele (Bashkirova and Lomvardas, 2019), thus the identification of this type of gene in an invariant T1-LAD follows expectations. Genes located in invariant T1-LADs were enriched for gene ontology (GO) terms that included detection of stimulus and sensory perception (Table 4). Another gene, *PAIP1*, located in an invariant T2-LAD, is a coactivator of the regulation of translation initiation (Fig. 5C). Genes located in invariant T2-LADs were enriched for GO terms that included various types of protein and proteolysis regulation, catabolic processes, and basic processes including vesicle, organelle, chromosome and membrane organization, a broad collection of terms that did not obviously showcase cell type specificity (Table 4). Collectively, these data showed that only a minority of the regions identified as T1-LADs or T2-LADs are conserved in their LAD subtype across all the cell types assayed, and that genes located within invariant LAD regions may be involved in universal functions or functions specific to a cell type absent from our atlas.

**Fig. 5:**
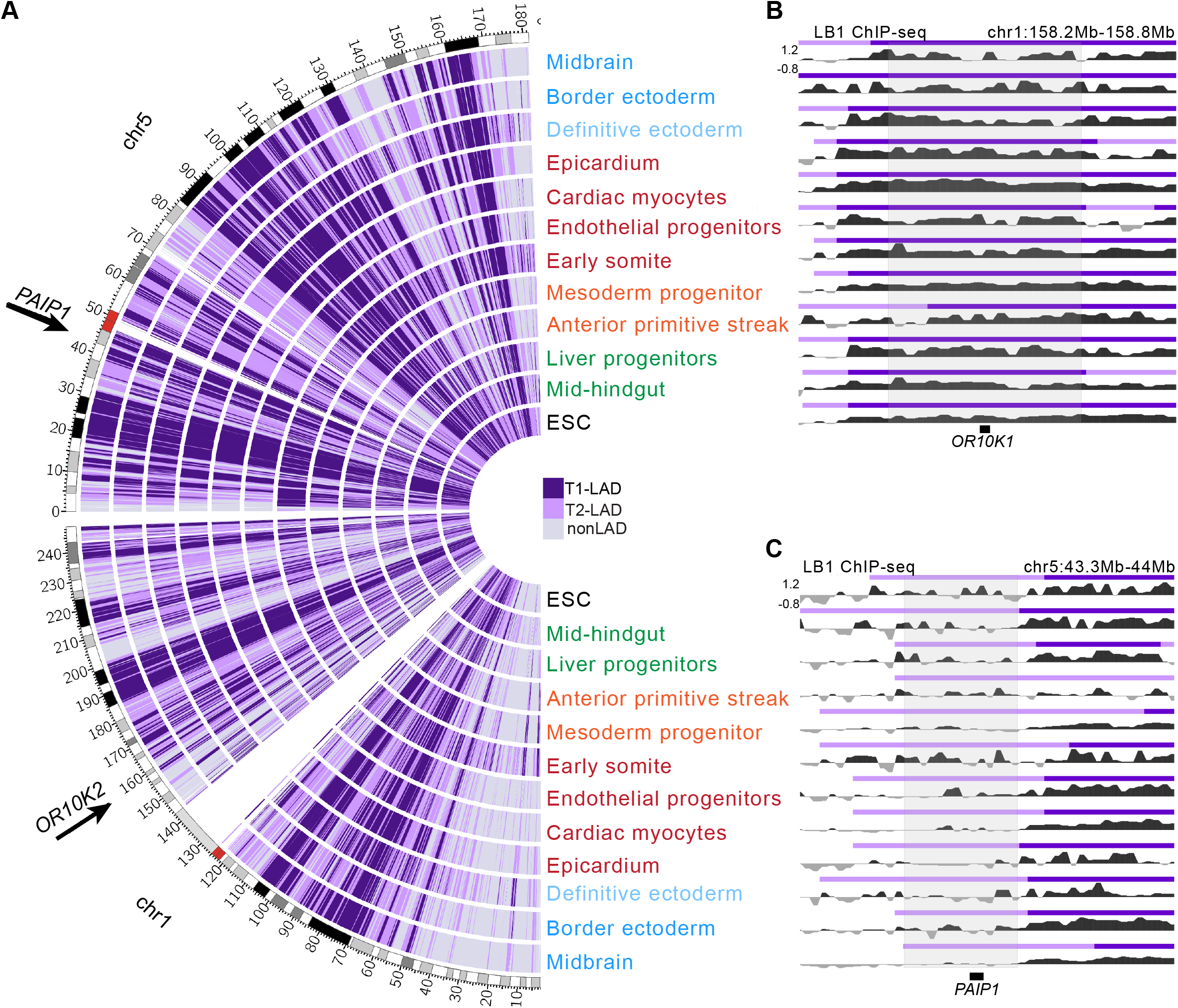
A subset of T1- and T2-LADs are invariant across cell types. A. LAD assignments across chromosomes 1 and 5 compared across the atlas. T1-LADs in dark purple, T2-LADs in light purple, nonLADs in grey. Arrows indicate example genes located in invariant T1- and T2- LADs. B. Genome browser LB1 tracks of the *OR10K1* locus, located in an invariant T1-LAD. C. Genome browser LB1 tracks of the *PAIP1* locus, located in an invariant T2-LAD. GO categories associated with B and C shown in Table 4. Y-axis indicated on the first track and consistent across all tracks shown for B and C.

### Integrating LAD states and gene expression across different cell types

Because we assayed 12 cell types and found relatively few invariant LADs, our atlas provides an opportunity to study LAD transitions and their association with gene regulation. In line with other studies showing cell type specific gene relocalization from LADs (Peric-Hupkes et al., 2010; Poleshko et al., 2017; Shah et al., 2021), we found that RNAseq expression changes between cell types generally corresponded to the expected change in LAD state (Fig. 6A). In several pairwise comparisons of ESCs to multiple cell types for which we had matched expression data, we found that genes which moved into a LAD subtype with increased LB1 occupancy were generally downregulated, while genes which moved to a subtype with decreased LB1 occupancy were generally upregulated (Fig. 6A). In accord, we found that genes that are canonical markers for various cell types are generally localized to nonLADs and highly expressed in the given cell type, although some reside in T2-LADs but are still highly expressed. Combined with the observation that these loci are not often found in T1-LADs, this suggests that T2-LADs may be an important transition for gene activation and expression compared to T1-LADs (Fig. 6B).

**Fig. 6:**
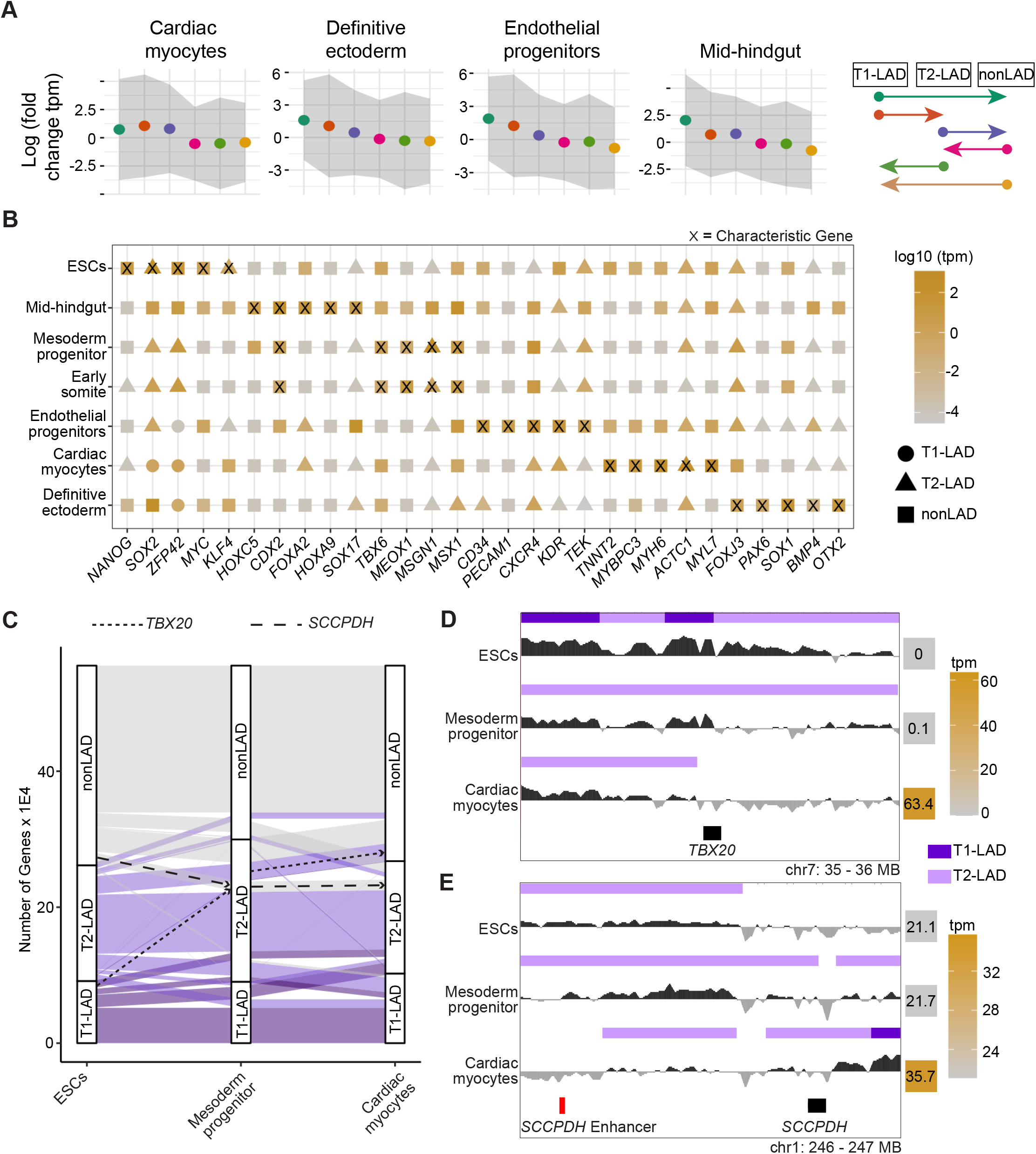
Gene expression changes broadly correspond with LAD states across developmental trajectories. A. Fold change in gene expression of differentiated cell types relative to ESCs for each LAD category change indicated by arrows in the panel on the right. Dots are mean and grey ribbons indicate standard deviation. ANOVA test is significant for all cell types. B. LAD assignments and expression levels of characteristic genes for each cell type show that many canonically expressed genes are nonLADs in their respective cell types and located within LADs in alternative cell types. C. Gene LAD assignment changes across cells from the mesoderm lineage. T1-LADs in dark purple, T2-LADs in light purple, nonLADs in grey. Genes rarely move from T1-LADs to nonLADs, with T2-LADs showing the greatest gene occupancy gains and losses. D,E: Track views of LB1 and LAD calls surrounding *TBX20* and *SCCPDH* and the *SCCPDH* enhancer show gene-LAD occupancy changes (*TBX20*) and enhancer-LAD occupancy changes (*SCCPDH* enhancer).

We assayed gene-LAD residency in cell types at various levels of lineage restriction of the mesoderm lineage, including mesoderm progenitors and more differentiated mesoderm-derived cell types, such as cardiac myocytes. We observed few genes that move from nonLAD to T1-LAD (or vice versa); instead, gene residency changes occur between T1- and T2-LADs or T2- and nonLADs (Fig. 6C, Supp. Fig. 6D), providing additional support for the idea that T2-LADs are an intermediate LAD subtype that may impact spatial positioning of biologically relevant loci during lineage restriction and gene activation. For example, *TBX20*, a gene integral to cardiac development (Kirk et al., 2007; Qian et al., 2008), moves from a T1-LAD in ESCs to a T2-LAD in mesoderm progenitors and to a nonLAD in cardiac myocytes where it becomes highly expressed, suggesting the involvement of lamina-mediated genome organization in transcriptional changes during development (Fig. 6D).

Some genes did not follow the expected shift in expression upon changing LAD type. For example, *SCCPDH*, which encodes an enzyme that exhibits oxidoreductase activity (Gogstad et al., 1981), shows increasing LB1 occupancy and an increase in gene expression across ESCs, mesoderm progenitors, and cardiac myocytes (Fig. 6E). It is possible that while the gene moved to a more repressive chromatin environment, its regulatory elements did not, which possibly facilitated the increased expression. We compared the locations of enhancers identified by genome-wide screens (Nasser et al., 2021) and observed that a cardiomyocyte enhancer of *SCCPDH* follows the opposite trend of its targeting gene, with decreased LB1 occupancy across the developmental trajectory, providing a potential explanation for the unexpected shift in expression of *SCCPDH* (Fig. 6E). Collectively, these results suggest that lamina-mediated organization of a gene, and potentially of its regulatory elements as well, may contribute to cell type-specific gene expression, and could provide an additional layer of transcriptional regulation. Moreover, we observed that organizational shifts in gene location appear to occur through T2- LADs and may occur in cell types earlier in the developmental trajectory where expression is not yet changed.

## Discussion

In this resource, we present genome-wide LB1 and H3K9me2 DNA binding across 12 cell types. The breadth of the datasets allows for LAD definitions and comparisons within and between multiple cell types, thereby providing additional clarity on the complex nature of lamina-associated chromatin. Work in *Drosophila* and murine cells identified subtypes of heterochromatin and/or LADs that share many characteristics with the LAD subtypes described in this work (van Bemmel et al., 2013; Filion et al., 2010; Zheng et al., 2015, 2018), and constitutive and facultative LADs have been identified on the basis of comparing between datasets. Our analyses revealed that lamina-associated chromatin is organized into at least two distinct types of LADs which are positioned distinctly. T1-LADs are a chromatin subtype that is gene poor, enriched in heterochromatin histone modifications, and generally inaccessible. T2-LADs are an intermediate, more accessible, more gene-rich chromatin subtype. Accordingly, gene expression in T2-LADs is generally higher than in T1-LADs. Our collective assessments suggest a model where distinct lamina-associated chromatin subtypes function and are maintained separately (Fig. 7).

**Fig. 7:**
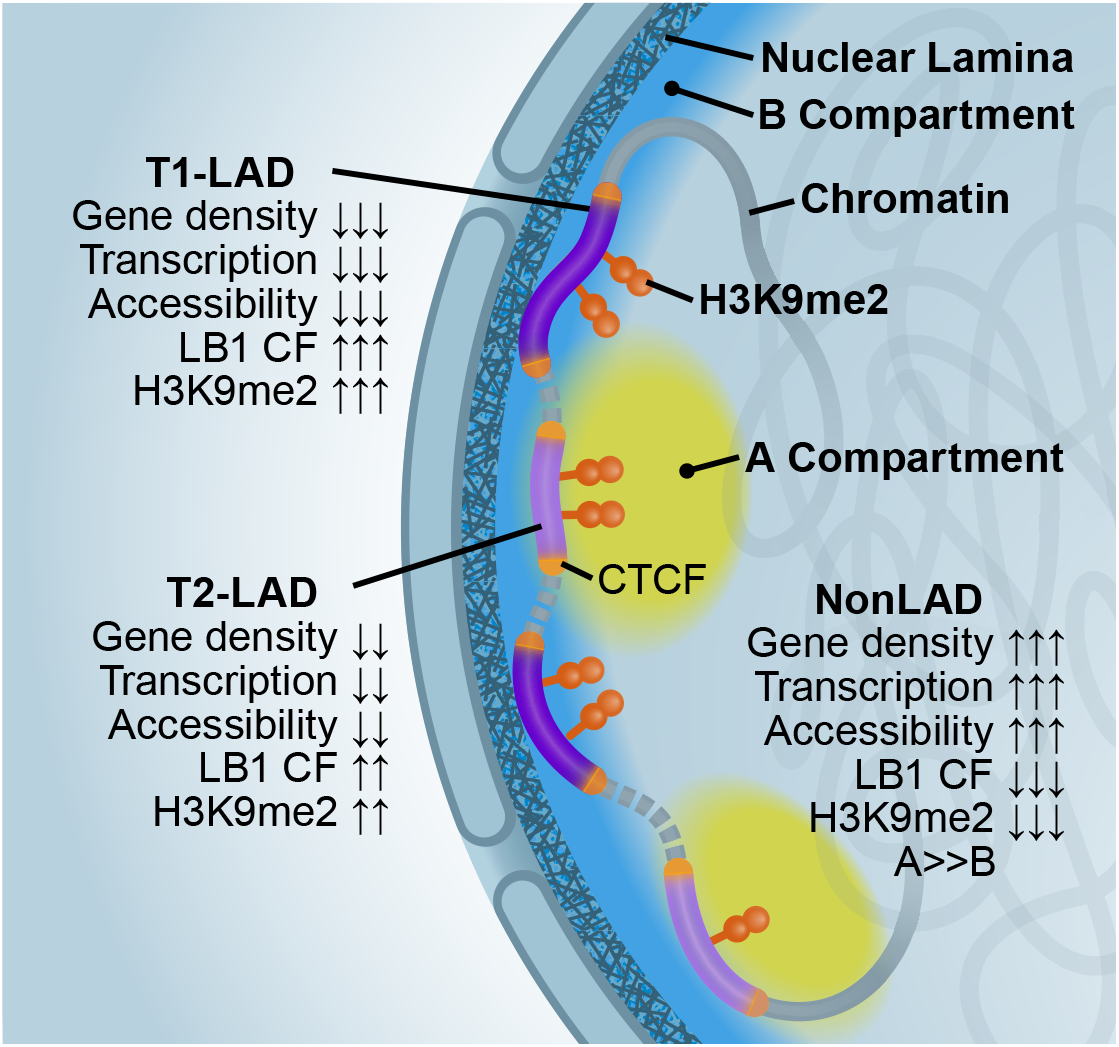
Graphical model of T1- and T2-LADs. Data from the atlas shows that T1-LADs have greater LB1 contact frequency (CF) and H3K9me2 enrichment, lower transcription, lower overall accessibility, lower gene density, and greater localization with the layer of peripheral LB1 compared to T2-LADs. This suggests that T2-LADs are a peripheral chromatin subtype localized to an intermediate layer of chromatin at the nuclear periphery that borders nonLADs.

Across all cell types analyzed in this work, we identified consistent characteristics that distinguish T1- from T2-LADs. One of the most striking is replication timing, which showed an enrichment for late domains in T1-LADs, compared to a depletion in T2-LADs. Other analyses showed that T1- LADs are the most inaccessible subtype, with the highest level of LB1 and H3K9me2 occupancy, strongest depletion of genes and accessible chromatin, and greatest enrichment of constitutively late replicating chromatin. In contrast, T2-LADs show lower levels of LB1 and H3K9me2 binding, less gene depletion and more chromatin accessibility, and are enriched for replication timing switch domains, raising the possibility of a more dynamic role for these regions. Consistently, recent studies revealed that facultative LADs, as defined by comparing LADs across four murine cell types, had lower LB contact frequency than constitutive LADs (Meuleman et al., 2013; Rooijers et al., 2019); however, our analyses identified different types of LADs without having to compare across cell types. Collectively, these results indicate that there exists an intermediate chromatin subtype between T1-LADs and nonLADs.

### LAD subtypes may be established and maintained by distinct mechanisms

Our FISH results in T2-LADs suggest that one possible mechanism driving the greater accessibility of T2- versus T1-LADs is consistent intermediate positioning of T2-LAD regions between the nuclear lamina chromatin and nonLAD chromatin, versus being embedded within the lamina, like T1-LADs. We do note, though, that T1-LADs show greater overlap with LB1 regions identified in single cell DamID data (Fig. 3D), supporting the possibility that a subset of T2-LADs may be more variably located within the nucleus. Future experiments assessing the frequency of T1- versus T2-LAD association with the nuclear lamina, coupled with single-cell assessments of transcriptional activity, such as single cell RNA FISH or combined single cell DamID and transcription assays (Rooijers et al., 2019), will greatly inform our understanding of the functional behavior that drives T1- and T2-LAD distinction, including the possibility that there may be T2-LAD subtypes where transcription is differentially regulated and/or localized to specific nuclear structures (Wang et al., 2021).

It is important to note that we identified LAD subtypes across a diverse series of cell types, but independent assessments of key LAD subtype features show strong consistency across cell types. Future studies aimed at understanding general mechanisms required for establishing or maintaining LADs, may reveal whether such mechanisms are distinct for T1- versus T2-LADs and if such differences drive localization to the lamina. Given the differences in transposable element enrichment, it will be exciting to determine if transposable elements contribute to differentiating T1 from T2-LADs. In *C. elegans*, molecular readers of H3K9 methylated chromatin are necessary to maintain localization of chromatin at the nuclear lamina (Gonzalez-Sandoval et al., 2015) and manipulation of H3K9me2 in murine cells results in abnormal lamina-associated chromatin organization (Poleshko et al., 2019). The differing levels of H3K9me2 occupancy we observed in T1-LADs compared to T2-LADs may play an important role in the establishment of these domains. Additional studies will be essential to understand how H3K9me2 contributes both to the complexity of chromatin subtypes at the nuclear periphery and how this histone modification may be required for normal positioning and/or repositioning of lamina-associated chromatin during development.

Our studies also indicated CTCF enrichment at the borders of T1- and T2-LADs, consistent with previous studies. Recent studies indicate that the boundary element CTCF and WAPL, a protein which facilitates unloading of cohesin, are likely to be important in the positioning of LADs with specific characteristics, including LAD size and LB1 density (Liu et al.; van Schaik et al.). We found these characteristics to be different between T1- and T2-LADs. In addition, since gene density is enriched in T2-LADs, normal positioning of a subset of T2-LADs flanked by CTCF and cohesin might be particularly sensitive to cohesin positioning factors, though additional experiments using degron based approaches in hiPSC-derived cell types will be required to directly test this prediction.

It is also interesting to consider that T1- and T2-LADs are associated with different replication timing domains, which have previously been associated with large-scale chromatin organization (Pope et al., 2014). These data collectively suggest that large scale chromatin compartments are likely more complex than a binary classification. Specifically, replication timing transition regions (genomic regions between early and late replicating domains), demarcating permissive versus repressive chromatin, exist between topologically associated domains, and show lamina-association (Johnstone et al., 2020). Here, we observed that T2-LADs, but not T1-LADs, share many of these transitional features, including higher gene density, higher basal transcription, and lower LB1 enrichment. This suggests that T2-LADs could be distinctly separated from T1-LADs by higher order chromatin organization, and play an active role in priming genes for transcription and enabling dynamic regulation of housekeeping genes. Given the relatively higher transcription in T2-LADs, another intriguing model is that a threshold level of transcription may be required to maintain heterochromatin in T2-LADs, compared to the particularly gene depleted and highly silenced T1-LADs (Denholtz et al., 2020; Grewal and Elgin, 2007).

### LADs are linked to cell type-specific transcription and are diverse amongst cell types

Maintenance of chromatin structure via association with the nuclear periphery contributes to gene regulation and is amongst the many factors contributing to cell identity and organogenesis (Meuleman et al., 2013; Peric-Hupkes et al., 2010; Poleshko et al., 2017; Robson et al., 2016). Gene expression signatures are correlated to changes in LAD occupancy, providing additional support for models in which LADs are critical regulators of cellular fate. Generally, canonical genes of particular developmental cell types showed an expected nonLAD-occupancy and high transcriptional signature. Across a developmental trajectory, genes that changed LAD state moved from nonLAD to T2-LAD or T2-LAD to T1-LAD, while few directly moved from nonLAD to T1-LAD (or vice versa), suggesting that the intermediate T2-LADs may be an important “priming” ground. Supporting that idea, poised promoters from chromHMM were enriched in T2-LADs). Additionally, we found examples of enhancers that gain and lose LAD occupancy in a cell type- and germline-specific manner, suggesting a role for LADs in cell type specification mediated by movement of regulatory elements towards and from the nuclear lamina.

Cell type specification and maintenance requires a complex stoichiometry of expression of many genes; perhaps the proportion of a cell population in which a locus is lamina-associated is yet another mechanism used to modulate levels of gene expression. As additional data are overlaid onto the foundational data of the atlas through future experiments, it will be exciting to identify a subset of LADs that may be linked to developmentally-relevant LAD changes. For example, recent data suggest LAMIN A/C filaments can act as transcriptional repressors (Buchwalter and Hetzer, 2017), and it is unclear if reduced lamin contact frequency of a particular locus at the nuclear periphery renders it susceptible to transcriptional activation and then re-localization, or vice versa. Our study has revealed several interesting candidates that can be used in future studies to reveal biological implications of T1- versus T2-LAD organization on cellular differentiation, perhaps in a hybrid single cell DNA- and RNA-FISH approach with or without super resolution microscopy.

The data in the atlas represents the largest compendium of LB1 and H3K9me2 ChIP-seq across non-cancer-derived human cell types to date. ChIP-seq provided several advantages essential for this work, including avoiding genetic manipulation of ESCs to insert the Dam-fusion protein, which could impact differentiation; circumventing challenges related to the different timescales for sufficient methylation to map LADs by DamID (Aughey et al., 2019; Rooijers et al., 2019; van Steensel and Henikoff, 2000) in cells with varied cell cycle dynamics; and allowing for assessment of H3K9me2, since a DamID fusion of a histone H3 gene would not distinguish the post-translational modification. We note that many seminal LAD studies have utilized DamID (Guelen et al., 2008; Robson et al., 2017; Vogel et al., 2007), which has distinct advantages (Aughey et al., 2019), particularly with regard to mapping chromatin-binding events in single cells. Such studies will be critical to determine the probability of T1- and T2-LAD lamina occupancy within a single cell, increasing our understanding of how distinct LAD subtypes behave and are established or maintained. Atlas cell types are derived from two genetic backgrounds that were highly replicable between each other and published reference datasets, but DamID in single cells could be used in the future to define potential small scale LB1 differences in polymorphic regions, which occur below the resolution of ChIP-seq. Ultimately, single cell DamID data could also be coupled with the atlas data to execute targeted studies of LAD signatures and organization. For example, atlas data can be used to design regions for super-resolution imaging, which may provide the necessary resolution to assess chromatin accessibility differences between subtypes at the nuclear periphery (Otterstrom et al., 2019; Ricci et al., 2015), which we observe by IF-FISH as differentially localized. This could be coupled with transcriptional assessments like RNA-FISH, thereby linking population-based peripheral chromatin organization to single cells in order to understand how organization contributes to transcription or vice versa and how this impacts normal development.

## Conclusion

This work reveals critical details about peripheral chromatin organization, specifically revealing consistent sub-segregation of peripheral chromatin across multiple human cell types and a cell type-specific set of overlapping but distinct lamin-associated chromatin domains. These findings lay the groundwork for future studies aimed at defining the driving cause behind the difference between T1- and T2-LADs. What is the physiologic consequence of having different subtypes of lamin-associated chromatin? How do overall peripheral chromatin organization and LAD/KDD subtypes impact genomic and cell type stability? Is this LAD segregation conserved in other species? The data resource described herein is a foundation upon which other data can be integrated to address these and other questions in the future. These investigations promise to expand our understanding of the role genome organization plays in establishing and maintaining cellular diversity, with broad potential impacts in health and disease.

## Methods

### Generation and maintenance of cell types

#### Human pluripotent stem cell maintenance

H9 human pluripotent stem cells were maintained in E8 media and passaged every four days onto matrigel-coated plates (Roche). ESCs, cardiomyocytes, epicardium, and endothelium were H9 ESC-derived. The remainder of the cells were H7 ESC-derived. H7 (WiCell) pluripotent stem cells were maintained in feeder-free conditions using mTeSR1 media (StemCell Technologies) + 1% penicillin/streptomycin (Thermo Fisher), and fresh media was added daily. Cells were cultured on tissue culture plastics coated with Geltrex basement matrix (Thermo Fisher; which was diluted 1:100 in DMEM/F12 media [Thermo Fisher] before being used to coat culture plastics). Prior to reaching confluence, H7 ESCs were dissociated using either Accutase (Thermo Fisher) or Versene (Thermo Fisher), and then were passaged onto new plates.

For all H7-derived cells, differentiation was conducted in serum-free media, either Chemically Defined Medium 2 (CDM2) or Chemically Defined Medium 3 (CDM3). The composition of CDM2 basal medium (Loh et al., 2014a, 2016) is: 50% IMDM + GlutaMAX (Thermo Fisher, 31980-097) + 50% F12 + GlutaMAX (Thermo Fisher, 31765-092) + 1 mg/mL polyvinyl alcohol (Sigma, P8136- 250G) + 1% v/v chemically defined lipid concentrate (Thermo Fisher, 11905-031) + 450 μM 1- thioglycerol (Sigma, M6145-100ML) + 0.7 μg/mL recombinant human insulin (Sigma, 11376497001) + 15 μg/mL human transferrin (Sigma, 10652202001) + 1% v/v penicillin/streptomycin (Thermo Fisher, 15070-063). Polyvinyl alcohol was brought into suspension by gentle warming and magnetic stirring, and the media was sterilely filtered (through a 0.22 μm filter) prior to use.

The composition of CDM3 basal medium (Ang et al., 2018) is: 45% IMDM + GlutaMAX (Thermo Fisher, 31980-097) + 45% F12 + GlutaMAX (Thermo Fisher, 31765-092) + 10% KnockOut serum replacement (Thermo Fisher, 10828028) + 1 mg/mL polyvinyl alcohol (Sigma, P8136-250G) + 1% v/v chemically defined lipid concentrate (Thermo Fisher, 11905-031) + 1% v/v penicillin/streptomycin (Thermo Fisher, 15070-063). Polyvinyl alcohol was brought into suspension by gentle warming and magnetic stirring, and the media was sterilely filtered (through a 0.22 μm filter) prior to use.

#### Cardiomyocyte differentiation

On day 0 (start of differentiation) human pluripotent stem cells were treated with 1mg/ml Collagenase B (Roche) for 1 hour, or until cells dissociated from plates, to generate embryoid bodies. Cells were collected and centrifuged at 300 rcf for 3 mins, and resuspended as small clusters of 50–100 cells by gentle pipetting in differentiation media containing RPMI (Gibco), 2 mM/L L-glutamine (Invitrogen), 4×10^4^ monothioglycerol (MTG, Sigma-Aldrich), 50 µg/ml ascorbic acid (Sigma-Aldrich). Differentiation media was supplemented with 2ng/ml BMP4 and 3 µmol Thiazovivin (Millipore). embryoid bodies were cultured in 6cm dishes (USA Scientific) at 37°C in 5% CO^2^, 5% O^2^, and 90% N^2^. On day 1, the media was changed to differentiation media supplemented with 30 ng/ml BMP4 (R&D Systems) and 30 ng/ml Activin A (R&D Systems), 5ng/ml bFGF (R&D Systems) and 1 µM Thiazovivin (Milipore). On day 3, embryoid bodies were harvested and washed once with DMEM (Gibco). Media was changed to differentiation media supplemented with 5 ng/ml VEGF (R&D Systems) and 5 µmol/L XAV (Stemgent). On day 5, media was changed to differentiation media supplemented with 5 ng/ml VEGF (R&D Systems). After day 8, media was changed every 3-4 days to differentiation media without supplements until approximately day 30.

#### Cardiomyocyte dissociation

Embryoid bodies were incubated overnight with 0.6mg/ml Collagenase Type II (Worthington) at 37°C. Dissociated cells were harvested and washed with Wash media (DMEM, 0.1% BSA) + 1 mg/ml DNase (VWR) twice and centrifuged at 300 rcf for 3 mins. Cells were resuspended in differentiation media supplemented with 1 µM Thiazovivin (Millipore) and filtered.

#### Epicardium differentiation

The human pluripotent stem cell cardiomyocyte protocol was followed up to day 3. On day 3, embryoid bodies were dissociated with TrypLE Express (Gibco). Dissociated cells were washed with Wash media (DMEM, 0.1% BSA) + 1mg/ml DNase (VWR) twice and centrifuged at 300 rcf for 3 mins. Cells were resuspended in differentiation media supplemented with 1ng/ml BMP4 (R&D Systems), filtered and counted using a hemocytometer. Cells were plated onto a matrigel-coated 96 well plate at 80,000 cells per well. On day 5, the media was changed to differentiation media supplemented with 5ng/ml VEGF (R&D Systems) and 1nM all-trans Retinoic Acid (Sigma-Aldrich). After day 5, media was changed every 2 days with the same day 5 differentiation media composition. On day 11, the media was changed to differentiation media supplemented with 5ng/ml VEGF (R&D Systems) and cells were fed with the same differentiation media every 2 days until day 15. On day 15, cells were dissociated with 1mg/ml Collagenase B (Roche) for 1 hour, washed with Wash media (DMEM, 0.1% BSA) + 1mg/ml DNase (VWR) and centrifuged at 300 rcf for 3 mins. Cells were further dissociated with 3ml TrypLE Express, washed with Wash media (DMEM, 0.1% BSA) + 1mg/ml DNase (VWR) and centrifuged at 300 rcf for 3 mins. Cells were resuspended in differentiation media supplemented with 1 µM Thiazovivin (Millipore), filtered and counted using a hemocytometer. Cells were plated in a matrigel-coated 6 well-plate at 100,000 cells per well. On day 16, media was changed to differentiation media supplemented with 5ng/ml VEGF (R&D Systems) and cells were fed every 2 days until they reached confluence (approximately Ddy 22). WT1 (WILMS-TUMOR 1) expression is indicative of successful epicardial differentiation (Zhou et al. 2008).

#### Endothelial cell differentiation

The human pluripotent stem cell cardiomyocyte protocol was followed up to day 5. At day 5, EBs were dissociated with TrypLE Express (Gibco). Dissociated cells were washed with Wash media (DMEM, 0.1% BSA) + 1mg/ml DNase (VWR) twice and centrifuged at 300 rcf for 3 mins. Cells were resuspended in differentiation media supplemented with 100ng/ml VEGF (R&D Systems) and 50ng/ml bFGF (R&D Systems), filtered and counted using a hemocytometer. Cells were plated onto a matrigel-coated 96 well plate at 80,000 cells per well. Media was changed every 2 days using differentiation media supplemented with 100ng/ml VEGF (R&D Systems) and 50ng/ml bFGF (R&D Systems). Cells were harvested and sorted on days 14-15.

#### Fluorescence-activated Cell Sorting (FACS)

Dissociated human pluripotent stem cell-derived cells were resuspended in differentiation media containing diluted antibodies (dilutions listed below) for 30 mins on ice. Cells were washed with differentiation media and resuspended in differentiation media + DAPI (1.35μg/ml, Biolegend) for FACS (BD FACSAria). Human pluripotent stem cell-derived cardiomyocytes used for IF-FISH were sorted by gating for SIRPA+ (PE-Cy7 anti-human CD172a/b, Biolegend, 1:200) and CD90- (APC anti-human CD90 (Thy1) Antibody, Biolegend, 1:200) cells. PSC-derived endothelial cells were sorted by gating for CD31+ (PE anti-human CD31, Biolegend, 1:200) cells.

#### Ectodermal differentiation

The day prior to beginning differentiation, H7 ESCs were dissociated with Accutase (Thermo Fisher) for 10 minutes at 37°C. Accutase was neutralized through the addition of excess DMEM/F12 media, and then ESCs were pelleted via centrifugation and the supernatant was aspirated. Pelleted ESCs were resuspended in mTeSR1 + 1% penicillin/streptomycin + 1 μM of the ROCK inhibitor Thiazovivin (Tocris) (henceforth referred to “cell-plating media”), and plated onto Geltrex-coated tissue culture plastics at a density of 4 x 10^5^ cells/cm^2^ (*i.e.*, 2.1 x 10^6^ cells per 10-cm dish). 24 hours after seeding, the cell-plating media was aspirated, and cells were briefly washed with DMEM/F12 to remove all traces of cell-plating media.

ESCs were then differentiated into definitive ectoderm through the addition of CDM2 basal medium supplemented with A8301 (1 μM; Tocris), LDN193189 (100 nM, Stemgent), C59 (1 μM; Tocris), and Trametinib (250 nM; Selleck Chem), which was added for 24 hours.

For border ectoderm induction, ESCs were differentiated into OTX2+ definitive ectoderm in 24 hours (as described above), and then definitive ectoderm was briefly washed (with DMEM/F12) and then treated with CDM2 basal media supplemented with BMP4 (30 ng/mL; R&D Systems), PD0325901 (100 nM; Tocris), CHIR99021 (3 μM; Tocris), and A8301 (1 μM; Tocris) for an additional 2 days, to generate, *PAX3+* border ectoderm progenitors. Differentiation media was aspirated and added fresh every 24 hours.

For midbrain induction, ESCs were differentiated into definitive ectoderm in 24 hours (as described above), and then definitive ectoderm was briefly washed (with DMEM/F12) and then treated with CDM2 basal media supplemented with A8301 (1 μM; Tocris), LDN193189 (100 nM; Stemgent), and Trametinib (250 nM; Selleck Chem) for an additional 2 days to generate neural progenitors. Neural progenitors were briefly washed (with DMEM/F12) and were then treated with CDM2 basal media supplemented with CHIR99021 (4 μM; Tocris), SB-505124 (2 μM; Tocris) and FGF2 (20 ng/mL; R&D Systems) for 2 days, to generate midbrain progenitors expressing *PAX2, PAX5, EN1, and EN2*. Differentiation media was aspirated and added fresh every 24 hours.

#### Endodermal differentiation

The day prior to beginning differentiation, H7 ESCs were dissociated with Accutase (Thermo Fisher) at 37°C. Accutase was neutralized through the addition of excess DMEM/F12 media, and then ESCs were pelleted via centrifugation and the supernatant was aspirated. Pelleted ESCs were resuspended in cell-plating media, and plated onto Geltrex-coated tissue culture plastics at a 1:8-1:16 cell seeding ratio. 24 hours after seeding, the cell-plating media was aspirated, and cells were briefly washed with DMEM/F12 to remove all traces of cell-plating media.

ESCs were then differentiated into anteriormost primitive streak through the addition of CDM2 basal medium supplemented with Activin A (100 ng/mL; R&D Systems), CHIR99021 (3 µM; Tocris), FGF2 (20 ng/mL; R&D Systems), and PI-103 (50 nM; Tocris), which was added for 24 hours. Day 1 anteriormost primitive streak cells were briefly washed (with DMEM/F12) and then differentiated into day 2 definitive endoderm through the addition of CDM2 basal medium supplemented with Activin A (100 ng/mL; R&D Systems), LDN-193189 (250 nM; Tocris), and PI-103 (50 nM; Tocris), which was added for 24 hours. Methods for anteriormost primitive streak and definitive endoderm formation have been described previously (Ang et al., 2018; Loh et al., 2014a, 2016; Martin et al., 2020).

For liver differentiation, day 2 definitive endoderm cells were briefly washed (with DMEM/F12) and further differentiated into day 3 posterior foregut through the addition of CDM3 base media supplemented with FGF2 (20 ng/mL; R&D Systems), BMP4 (30 ng/mL; R&D Systems), TTNPB (75 nM; Tocris), A8301 (1 μM; Tocris). Day 3 posterior foregut cells were briefly washed (with DMEM/F12), and then further differentiated on days 4–6 with CDM3 base media supplemented with Activin A (10 ng/mL; R&D Systems), BMP4 (30 ng/mL; R&D Systems), Forskolin (1 μM; Tocris) to generate liver bud progenitors expressing *HNF4A* and *TBX3*. Methods for liver bud progenitor formation have been described previously (Ang et al., 2018; Loh et al., 2019; Martin et al., 2020).

For mid-hindgut differentiation, day 2 definitive endoderm cells were briefly washed (with DMEM/F12) and further differentiated into day 6 mid-hindgut progenitors expressing *FOXA2*, *CDX2* and *HOXA9* through the addition of CDM2 base media supplemented with FGF2 (100 ng/mL), BMP4 (10 ng/mL) and CHIR99021 (3 μM) for 4 days. Methods for mid-hindgut progenitor formation have been described previously (Ang et al., 2018; Loh et al., 2014a).

#### Mesodermal differentiation

The day prior to beginning differentiation, H7 ESCs were dissociated with Accutase (Thermo Fisher) at 37°C. Accutase was neutralized through the addition of excess DMEM/F12 media, and then ESCs were pelleted via centrifugation and the supernatant was aspirated. Pelleted ESCs were resuspended in cell-plating media, and plated onto Geltrex-coated tissue culture plastics at a 1:8-1:16 cell seeding ratio. 24 hours after seeding, the cell-plating media was aspirated, and cells were briefly washed with DMEM/F12 to remove all traces of cell-plating media.

ESCs were then sequentially differentiated into anterior primitive streak, mesoderm progenitors (enriched in *TBX6*, *CDX2*, and *MSGN1* expression) and early somites (enriched in *MEOX1* expression) as described previously (Loh et al., 2016). Briefly, ESCs were differentiated into anterior primitive streak through the addition of CDM2 basal medium supplemented with Activin A (30 ng/mL; R&D Systems), CHIR99021 (4 μM; Tocris), FGF2 (20 ng/mL; R&D Systems), and PIK90 (100 nM; Calbiochem), which was added for 24 hours, thus generating day 1 anterior primitive streak (Loh et al., 2016).

For mesoderm induction, ESCs were differentiated into anterior primitive streak in 24 hours (as described above), and then anterior primitive streak was briefly washed (with DMEM/F12) and then treated with CDM2 basal media supplemented with A8301 (1 μM; Tocris), LDN193189 (250 nM; Tocris), CHIR99021 (3 μM, Tocris), and FGF2 (20 ng/mL; R&D Systems), which was added for 24 hours, thus generating mesoderm progenitors (Loh et al., 2016).

For early somite induction, ESCs were differentiated into anterior primitive streak and then further differentiated into mesoderm (as described above). Mesoderm was briefly washed (with DMEM/F12) and then treated with CDM2 basal media supplemented with CDM2 base media supplemented with A8301 (1 μM; Tocris), LDN193189 (250 nM; Tocris), XAV939 (1 μM; Tocris), and PD0325901 (500 nM; Tocris) for 24 hours, thus generating early somite progenitors(Loh et al., 2016).

ESCs were differentiated into primitive streak through the addition of CDM2 basal medium supplemented with Activin A (30 ng/mL; R&D Systems), BMP4 (40 ng/mL; R&D Systems), CHIR99021 (6 μM; Tocris), FGF2 (20 ng/mL; Thermo Fisher) for 24 hours (with the optional addition of PIK90 (100 nM; Calbiochem)), as previously described (Loh et al., 2016).

### IF-FISH, Imaging, and Quantification

#### IF and IF-FISH

ESCs and cardiac myocytes were grown and/or differentiated in culture, sorted (see above) and plated for FISH by direct growth on coverslips. Cells were fixed with 4% paraformaldehyde (PFA) for 10 minutes at room temperature (RT), and permeabilized with 0.5% Triton X-100 for 10 minutes at RT. Permeabilized cells were then blocked in 1% BSA in PBS-T (8mM Na2HPO4, 150mM NaCl, 2mM KH2PO4, 3mM KCl, 0.05% Tween 20, pH 7.4) and incubated with primary and secondary antibodies for 1 hour each at RT with 3 PBS-T washes for 5 minutes each in between antibody incubations. Primary antibodies used were anti-Lamin B1 (1:1000, Abcam #ab16048) and anti-H3K9me2 (1:1000, Active Motif #39239). Secondary antibodies used were anti-rabbit AlexaFluor 488 (1:1000, Invitrogen #21206), anti-rabbit AlexaFluor 568 (1:1000, Invitrogen #10042).

#### IF-FISH

Following IF, cells were post-fixed with 2% PFA for 10 minutes at RT and permeabilized with 0.7% Triton X-100 for 10 minutes at RT. Cells were incubated in 2x SSC-T (3.0M NaCl, 0.3M Sodium Citrate, 0.1% Tween 20) for 5 minutes at RT, followed by washes in 2x SSC-T with 50% formamide for 5 minutes at RT, 2.5 minutes at 92°C, and 20 minutes at 60°C. Cells were hybridized with a Cy2, Cy3, or Cy5 directly-labeled DNA probe diluted in a hybridization mix containing 50% formamide, 1x dextran sulfate sodium salt (Fisher Scientific #BP1585) with PVSA (Poly(vinylsulfonic acid, sodium salt) solution (Sigma #278424), 10μg RNAseA, 10mM dNTPs, and 2-5pmol probe for 30 minutes at 80°C, then overnight (minimum 16 hours) at 37°C. Probes were designed in target chromosomal regions (see Table 5) using a tiled oligo-based approach with 80-mer probes spaced at 4 probes/kb in designated chromosomal regions. Cells were washed with 2x SSC-T at 60°C for 15 minutes followed by washes in 2x SSC-T and 2x SSC for 10 minutes each at RT. Cells were counterstained with DAPI solution (Sigma #D9542) diluted in 2x SSC for 5 min at RT. Cells were mounted on coverglass with SlowFade Gold antifade mounting reagent (Invitrogen #S36936) prior to image acquisition. The nonLAD probe was not dye-conjugated; for this probe, IF-FISH was performed as above with the following modifications. Primary hybridization mix contained 5μg RNAseA, 5mM dNTPs, and a probe concentration of 50pmol. Following primary probe hybridization, cells were washed with 2x SSC=T at 60°C for 15 minutes, followed by 2x SSC-T and 2x SSC for 10 minutes. A second hybridization mix was added containing 10% formamide, 1x dextran sulfate salt with PVSA, and 10pmol of a secondary probe conjugated to AlexaFluor 647. Cells were hybridized for 2 hours at RT, then washed with 2x SSC-T at 60°C for 10 minutes, then 2x SSC-T, and 2x SSC for 5 minutes each. Samples were then counterstained with DAPI and mounted, as above.

#### Imaging

Confocal 3D images were taken using 120nm step Z-stacks, with an approximate range of 10-70 Z-planes per cell. Obtained images were deconvoluted using Leica Lightning Deconvolution Software. Representative images in Fig. 4 represent a single focal plane with brightness and contrast adjusted equivalently across samples in ImageJ. 3D reconstructions were performed using IMARIS v.7.4 software (Bitplane AG, Switzerland). Nuclear lamina and H3K9me2 surfaces were created using Surfaces tool with automatic settings based on the fluorescent signals from the anti-LB1 and anti-H3K9me2 antibodies. DNA FISH foci were generated using the Spots tool with a 300nm diameter, created at the intensity mass center of the fluorescent probe signal. Distance from the center of the FISH focus to the inner and outer edge of the nuclear lamina surface was quantified using the Measurement Points tool. Thickness of the H3K9me2-marked peripheral heterochromatin layer was calculated as the distance between the inner and outer edge of the H3K9me2 surface quantified using the Measurement Points tool with 5 random measurements per cell, from 10 independent cells per cell type (total of 50 measurements per cell types). For ESCs and cardiac myocytes, the 90% percentile of H3K9me2 thickness was measured at 349nm and 422nm, respectively. In cases when the signal from a FISH focus was embedded into the nuclear lamina layer, the measurement returned negative distances. Statistical significance of the differences in localization between T1-, T2-, and nonLAD foci (Fig. 4A and 4B) was calculated by a Kruskal-Wallis test with post-hoc Dunn test for multiple comparisons. Significance measurements are provided in figure legends. To compare the localization of the additional probes tested (Fig. 4C) to the matched LAD type probe in Fig. 4A and 4B, a Mann Whitney U test was performed; no significant differences were observed in T1- to T1-LAD or T2- to T2-LAD probe comparisons in either ESCs or CMs.

### ChIP, ChIP-qPCR, and ChIP-seq Library Preparation

#### ChIP

Undifferentiated ESCs and all differentiated cell types were crosslinked in culture by addition of methanol-free formaldehyde (ThermoFisher, final 1% v/v), and incubated at room temperature for 10 minutes with gentle rotation. Crosslinking was quenched by addition of glycine (final 125mM) and incubated at room temperature for 5 minutes with gentle rotation. Media was discarded and replaced with PBS; cells were scraped and transferred to conical tubes and pelleted by centrifugation (250xg, 5 minutes at room temperature). Resulting pellets were flash frozen on dry ice and stored at −80C. For ChIP, 30uL protein G magnetic beads (per ChIP sample; Dynal) were washed 3 times in blocking buffer (0.5% BSA in PBS); beads were resuspended in 250uL blocking buffer and 2ug antibody (LaminB1: ab16048 [abcam]; H3K9me2: ab1220 [Abcam]) and rotated at 4C for at least 6 hours. Crude nuclei were isolated from frozen crosslinked cells as follows: cell pellet (from 10cm plate) was resuspended in 10mL cold Lysis Buffer 1 (50mM HEPES-KOH pH7.5, 140mM NaCl, 1mM EDTA, 10% Glycerol, 0.5% NP-40, 0.25% Triton X-100, and protease inhibitors), and rotated at 4C for 10 minutes, followed by centrifugation (250xg, 5 minutes at room temperature). Supernatant was discarded and the pellet was resuspended in 10mL cold Lysis Buffer 2 (10mM Tris-HCl pH 8.0, 200mM NaCl, 1mM EDTA, 0.5mM EGTA, and protease inhibitors), and rotated at room temperature for 10 minutes, followed by centrifugation (250xg, 5 minutes at room temperature). Supernatant was discarded and nuclei were resuspended/lysed in 1mL cold Lysis Buffer 3 (10mM Tris-HCl, pH 8.0, 100mM NaCl, 1mM EDTA, 0.5mM EGTA, 0.1% Na-Deoxycholate, and protease inhibitors) and transferred to pre-chilled 1mL Covaris AFA tubes (Covaris). Samples were sonicated using a Covaris S220 sonicator (high cell chromatin shearing for 15 minutes; Covaris). Lysates were transferred to tubes and Triton X-100 was added (final 1%) followed by centrifugation (top speed, 10 minutes at 4C in microcentrifuge). Supernatant was transferred to a new tube; protein concentration was measured by Bradford assay. Antibody-conjugated beads were washed 3 times in blocking buffer, resuspended in 50uL blocking buffer and added to 500ug input protein for overnight incubation with rotation at 4C. 50ug lysate was aliquoted and stored at −20C for input. On day 2, beads were washed 5 times in 1mL RIPA buffer (50mM HEPES-KOH pH 7.5, 500mM LiCl, 1mM EDTA, 1% NP-40, 0.7% Na-Deoxycholate) with 2-minute incubation at room temperature with rotation for each wash. Beads were washed in 1mL final wash buffer (1xTE, 50mM NaCl) for 2 minutes with rotation at room temperature before final resuspension in 210uL elution buffer (50mM Tris-HCl pH 8.0, 10mM EDTA, 1% SDS). To elute, beads were incubated with agitation at 65C for 30 minutes. 200uL eluate was removed to a fresh tube, and all samples (ChIP and reserved inputs) were reverse-crosslinked overnight at 65C with agitation for a minimum of 12 hours, but not more than 18 hours. 200uL 1xTE was added to reverse crosslinked DNA to dilute SDS, and samples were RNaseA treated (final 0.2mg/mL RNase; 37C for 2 hours) and Proteinase K (final 0.2mg/mL Proteinase K; 55C for 2 hours) before phenol:chloroform extraction and resuspension in 10mM Tris-HCl pH 8.0. ChIP and input DNA was quantified by Qubit (ThermoFisher).

#### ChIP-qPCR

Post quantification, ChIP DNA from ESCs was diluted 1:5 and used for qPCR assessment across 20 independent T1-LAD, T2-LAD, nonLAD and T1-KDD, T2-KDD, and nonKDD regions (primer sequences in Table 6). qPCR was performed in 10uL reactions in 384- well format with 2uL 1:5 diluted template, 2x Power SyBr mastermix (Thermo Fisher) and 0.1uM each forward and reverse primer. qPCR reactions were run for 40 cycles using standard conditions [3 minutes at 95°C; 40x (15 seconds at 95°C; 1 minute at 60°C)] on a QuantStudio 5 or QuantStudio 7 qPCR machine (Applied Biosystems). For qPCR assessments, average enrichment (average Ct ChIP/average Ct input) were quantified per primer set.

#### Library Preparation

ChIP-seq libraries were prepared using the NEBNext Ultra II DNA library prep kit (NEB). Samples were indexed for multiplex sequencing. Library quality was analyzed by BioAnalyzer (Agilent Genomics) and quantified using qPCR (Kapa Biosystems or NEB). Libraries were pooled for multiplex sequencing, re-quantified, and sequenced on the Illumina NextSeq500 platform (vII; 75bp single-end sequencing; Illumina).

### RNA Isolation and RNA-seq Library Preparation

Undifferentiated ESCs or differentiated cells were scraped from tissue culture plates with 1xPBS, and centrifuged at 1500g for 5 minutes at room temperature. After discarding supernatant, cell pellets were flash frozen in dry ice and stored at −80C until processing. RNA was isolated using QIAGEN RNeasy total RNA extraction kit (QIAGEN). RNA quality was analyzed by BioAnalyzer; samples with RIN scores >8 were chosen for further processing. RNA libraries were prepared using the NEBNext Ultra II DNA Library Prep kit (NEB) with the NEBNext Poly(A) mRNA Magnetic Isolation Module (NEB) to enrich for poly-A tailed RNA molecules. RNA-seq library quality was analyzed by BioAnalyzer (Agilent Genomics) and quantified using qPCR (Kapa Biosystems). Libraries were pooled for multiplex sequencing, re-quantified, and sequenced on the Illumina NextSeq500 platform (vII; 75bp single end sequencing; Illumina).

### ChIP-seq/RNA-seq Processing and Computational Analyses

#### ChIP-sequencing data processing for LAMIN B1 and H3K9me2

Adapters were trimmed using Trimmomatic [v0.39] (Bolger et al., 2014). Sequencing reads were aligned to human reference hg38 using BWA-MEM [v0.7.17] (Li and Durbin, 2010). Aligned reads were converted to BAM and sorted using Samtools [v0.1.19] (Li et al., 2009), with quality filter (“-F”) set to 1804. Duplicates were removed using Picard [v2.18.7] MarkDuplicates. Sequencing reads from the ENCODE blacklist were removed using Bedtools [v2.29.0] (Li et al., 2009; Quinlan and Hall, 2010). Two biological replicates were analyzed for each cell type. The data for cell types based on combined replicates adhere to ENCODE3 standards (Table 1) (ENCODE Project Consortium et al., 2020).

#### Identification of LADs and KDDs

LB1 and H3K9me2 ChIP-seq signals were calculated and converted into BedGraph files using deepTools bamCompare [v3.3.2] (Ramírez et al., 2014) with 20kb bins, using the signal extraction scaling method (Diaz et al., 2012) for sample scaling, followed by quantile normalization between cell types to decrease the impact of batch effects. The bin size of 20kb was chosen based on assessment of the literature and motivation to describe LADs in as fine of resolution as possible (Borsos et al., 2019; Kind et al., 2015; Zheng et al., 2018). HMMs were implemented for each cell type using pomegranate [v0.11.1] (Akaike, 1974; Schreiber, 2017). Each HMM was initialized using a normal distribution and k-means with a uniform transition matrix, and trained using the Baum-Welch algorithm. Each cell type-specific model was then applied to predict LAD or KDD state genome-wide per 20kb bin, using the median value from both replicates for each bin, for each cell type individually, filtering regions in the ENCODE blacklist from consideration. For the LAD predictions, states were labeled as T1-LAD, T2-LAD or nonLAD based on median LB1 signal for the bins with that state label, with the highest median LB1 signal being assigned T1-LAD, second highest T2-LAD, and lowest nonLAD. The same strategy was employed to assign T1-, T2- and nonKDDs.

#### LAD boundary analyses

LB1 occupancy at LAD boundaries were computed using the computeMatrix --referencePoint and plotProfile tools from deepTools [v3.3.2] (Ramírez et al., 2014).

#### RNA-sequencing analysis

Transcriptome data were quantified using Kallisto [v0.44.0] quant with fragment length determined by BioAnalyzer, standard deviation of 10, and 30 bootstraps, assigning reads using the Ensembl [v96] genome annotation (Bray et al., 2016). TPM values were quantile-normalized between cell types. Differentially expressed transcripts (q<=0.01) between cell types were identified using Sleuth [0.30.0] (Pimentel et al., 2017).

#### ATAC-seq analysis

ATAC peaks from H9-derived cells (Liu et al., 2017) were downloaded as BED files from GEO and lifted over from hg19 to hg38, taking the intersection of two replicates for each cell type.

#### CTCF analysis

Bigwig files were downloaded from GEO (see Data Access section). CrossMap was used to lift over bigwigs from hg19 to hg38.

#### Hi-C analysis

Hi-C data for CMs and ESCs were downloaded as Cooler files from the 4D Nucleome Data Portal (Zhang et al., 2019). A and B compartments were called using cooltools [v0.3.0] (Abdennur and Mirny, 2020).

#### Enrichment analyses

Odds ratios were calculated based on two by two tables of counts of 20kb genomic bins for category (T1-LAD, T2-LAD or KDD) overlap and domain of interest (replicating timing domain, gene, transposable element, etc.) overlap. P-values were calculated by Fisher’s exact test.

#### Comparison with single cell DamID

Single cell DamID data from 172 KBM7 cells from clone 5-5 was downloaded from the Gene Expression Omnibus (GSE68260) in hg19 and lifted over using pybedtools to hg38, removing regions that did not lift over. These data were intersected with T1- and T2-LADs using pybedtools (Dale et al., 2011).

#### Gene ontology enrichment analyses

Enriched gene ontology terms for genes located in invariants T1- or T2-LADs was done using the HumanBase Modules tool against a background set of all genes.

#### Supporting analyses

Gene annotations used throughout are from Ensembl v96. The reference genome used was human hg38, downloaded from the UCSC Genome Browser. Constitutive late, constitutive early and switch domains were obtained from (Dixon et al., 2018). They defined replication timing domains by their consistency across multiple cell types, thus the same domains are used in each cell type in our analysis. Transposable elements from RepeatMasker were downloaded from the UCSC Genome Browser. Plotting, statistical analyses and supporting analyses were conducted in Python [v3.6] with packages Jupyter, matplotlib (Hunter, 2007), seaborn (Waskom, 2021), upsetplot (Waskom, 2021), scikit-learn (Pedregosa et al., 2011), numpy (der Walt et al., 2011), pybedtools (Dale et al., 2011; Quinlan and Hall, 2010), Circos (Krzywinski et al., 2009) and deepTools [v3.3.2] (Ramírez et al., 2014) and in R [v4.1.0] (Computing and Others, 2013) with packages dplyr (Wickham et al., 2015), tidyverse (Wickham et al., 2019), and ggalluvial (Brunson, 2020).

#### Violin and box plots

Boxes represent standard median (center dot or line) and interquartile range (25th to 75th percentile). Whiskers denote 1.5x interquartile range.

### Data Access for Data Generated in this Paper

RNAseq data from ventricular CMs, ESCs and endothelial cells, ChIP-seq data for LB1 and H3K9me2 from all cell types, LAD and KDD calls are available through the Gene Expression Omnibus (GSE155244).

### Data Access for Previously Published Data

#### RNA-seq

Data from day 3 early somite and mesoderm progenitor cells were downloaded from SRP073808 (Koh et al., 2016). Data from mid-hindgut were downloaded from SRP033267 (Loh et al., 2014). Data from neural ectoderm were downloaded from SRP113027 (Tchieu et al., 2017).

#### ATAC-seq

Data from ESCs, CMs and day 2 mesoderm were downloaded from GSE85330 (Liu et al., 2017).

#### CTCF ChIP-seq

Data from ESCs were downloaded from GSM3263064 and GSM3263065, from day 2 mesoderm from GSM3263066 and GSM3263067 and from day 80 ventricular CMs from GSM3263074 (Zhang et al., 2019).

#### Hi-C

Data from ESCs were downloaded from GSM3263064 and GSM3263065 and from day 80 ventricular CMs from GSM3263074 (Zhang et al., 2019).

#### Transcription factor binding sites

Transcription factor clustered binding sites from the ENCODE project were downloaded from the UCSC Genome Browser (ENCODE Project Consortium, 2012; ENCODE Project Consortium et al., 2020).

#### chromHMM states

chromHMM states in reference genome hg38 from the 25 state model were downloaded for cell types E008, E095, E066, and E013 as the closest cell types matching ESCs, cardiac myocytes, liver cell, and mesoderm progenitor cells from https://egg2.wustl.edu/roadmap/web_portal/imputed.html.

### Code Availability

Code used to train the HMMs described herein and for the main figures is available at https://github.com/keoughkath/LAD_atlas.

## Supporting information

KeoughandShah_Table5

KeoughandShah_Table1

KeoughandShah_Table2

KeoughandShah_Table3

KeoughandShah_Table4

KeoughandShah_Table6

KeoughandShah_SuppFigs

## Acknowledgements

We thank Geoffrey Fudenberg, Vijay Ramani, Abigail Buchwalter, Ashley Karnay, Wonho Kim, Andrey Poleshko, Cheryl Smith, Jonathan A. Epstein, and Benoit Bruneau for valuable conversations and critical review of the manuscript. We thank Jonas Fowler, Xiaochen Xiong, and Alana Nguyen for assistance with validating differentiated cells. We thank Kathryn Claiborn for copy editing this manuscript, and Giovanni Maki for help making the model figure. K.C.K. was supported in part by a Discovery Fellowship. K.C.K. and K.S.P. were supported by NHLBI (HL098179), the NIH 4D Nucleome program (GM140324), and Gladstone Institutes. C.E.D. was supported by the Ford Foundation Predoctoral Fellowship and the Stanford Graduate Fellowship. A.C. and R.E.A.S. were supported by the California Institute for Regenerative Medicine (Bridges Program TB1-01195). K.M.L. is a Packard Foundation Fellow, Pew Scholar, Human Frontier Science Program Young Investigator, Baxter Foundation Faculty Scholar and The Anthony DiGenova Endowed Faculty Scholar. R.J was supported by a Burroughs Wellcome Career Award for Medical Scientists, NSF CMMI-1548571, funds from the American Heart Association and Allen foundation, and a NIH New Innovator Award (DP2, HL147123).

## Disclosure Declaration

K.K. is currently an employee of FaunaBio. A.C. is currently an employee of Orca Bio. K.S.P. is a consultant for Tenaya Therapeutics.

## Abbreviations

ChIP: chromatin immunoprecipitation
ESC: embryonic stem cell
FISH: fluorescence *in situ* hybridization
GO: gene ontology
H3K9me2: dimethylated histone H3 lysine 9
HMM: Hidden Markov Model
IQR: interquartile range
kb: kilobases (1,000 basepairs)
KDD: K9-dimethyl domain, or H3K9me2-associated domain
LAD: lamina-associated domain
LB1: LAMIN B1
Mb: megabases (1,000,000 basepairs)
qRT-PCR: quantitative reverse transcription PCR
T1-KDD: Type 1 KDD
T1-LAD: Type 1 LAD
T2-KDD: Type 2 KDD
T2-LAD: Type 2 LAD
TPM: transcripts per million reads

---

**Supp. Fig. 1: Validation of hESC-derived cell types.** A-F. Genes as indicated. Panels in (A, C) (left), and (E) (left) show downregulation of pluripotency genes as cells differentiate into definitive ectoderm, border ectoderm, and mid-hindgut. All other panels show enrichment of lineage-specific genes, as indicated. For all assessments, relative enrichment is shown (normalized to hESCs). n=3 biological replicates. Error bars=SEM. G. Schematic of epicardial specification. H. WT1 and DAPI staining of cardiac cells (left) and epicardial cells (right). I. Quantification of WT1 positive and negative cells from (H). J. Schematic of endothelial specification. K. Endothelial cell sorting using FSC and CD31 markers. L. Schematic, cardiac differentiation. M. Cardiac cells show canonical TNNT2 expression.

**Supp. Fig. 2: LB1 antibody ChIP and specificity validation.** A. LB1 ChIP-immunoblot of ESCs shows enrichment of LB1 in LB1 ChIP (lane 1) compared to no antibody and IgG controls (lanes 3 and 4, respectively). B. LB1 antibody specificity measured by ChIP immunoblot in mesoderm progenitors and mid-hindgut cell lysates with and without peptide competition shows LB1 enrichment in ChIP samples without peptide (lanes 1 and 2), but not in ChIP with LB1 peptide (lane 3).

**Supp. Fig. 3: Validation of LADs identified by a 3-state HMM.** A. AIC differences between numbers of LAD states. B. BIC differences between numbers of LAD states. C. LB1 enrichment measured by ChIP-qPCR in three LAD states shows greatest LB1 enrichment in T1-LADs. Each box represents ChIP/input average of three independent ChIP-qPCR assessments at 20 loci. D. Overlap of LADs from ESCs identified in this work with previously published LADs identified with DamID (Meuleman et al. 2013).

**Supp. Fig. 4: H3K9me2 antibody ChIP and specificity validation.** A. H3K9me2 ChIP-immunoblot of ESCs shows enrichment of H3K9me2 in H3K9me2 ChIP (lane 3) compared to input and no antibody controls (lanes 1 and 3, respectively). B. H3K9me2 antibody specificity measured by dot blot shows positive signal in total ESC cell lysate (lane 1, dots 1 and 2) and with H3K9me2 peptide (lane 1, dot 4), but not H3K9me1 or H3K9me3 (lane 1, dots 1 and 3, respectively). No antibody binding detected in buffer only controls (lane 2).

**Supp. Fig. 5: H3K9me2 HMM validation; KDDs and LADs are highly overlapping across the atlas.** A. AIC differences between numbers of KDD states. B. BIC differences between numbers of KDD states. C. H3K9me2 binding in KDDs for each cell types. D. H3K9me2 enrichment measured by ChIP-qPCR in three KDD states. E. Example track of a genomic region with overlapping LADs and KDDs in indicated cell types. F. Overlap between T1-LADs and T1-KDDs, and T2-LADs and T2-KDDs across the Atlas datasets show a high degree of overlap. G. H3K9me2 occupancy in LADs for all atlas cell types.

**Supp. Fig. 6: H3K9me2 IF and cell sorting for IF-FISH; T2-LADs showcase the highest gene movement across developmental trajectories.** A. Representative image (individual slices of Z-stacks, cropped for maximal visualization) of H3K9me2 IF in both ESCs (top) and sorted cardiac myocytes (bottom). Quantification of the localization of the H3K9me2 signal to the nuclear periphery (5 independent measurements in n=10 nuclei per cell type; see Methods). Scale bars = 1µM. B. Fluorescence activated cell sorting for pure cardiac myocyte populations via SIRPA, used for IF-FISH. C. Probe tracks (left), representative images (middle) and quantification (right) of an additional T1-LAD in cardiac myocytes. Black bar in quantification plot marks the median. D. Gene LAD assignment changes across cells from the mesoderm lineage, as indicated. T1- LADs in dark purple, T2-LADs in light purple, nonLADs in grey. Genes rarely move from T1-LADs to nonLADs, with T2-LADs showing the greatest gene occupancy gains and losses.

**Table 1: Sequencing statistics for LAD atlas datasets.** Total number of unique reads sequenced per ChIP condition (LB1, H3K9me2 or Input) per replicate in all atlas cell types. Replicability measure (Pearson correlation) scores provided for LB1 and H3K9me2 replicate sets.

**Table 2: HMM-identified LADs in the LAD atlas.** T1- and T2-LADs for each cell type in the atlas in .bed format. Cell types separated by worksheets. All coordinates are in hg38. Each sheet can be exported into a .bed file and uploaded to a genome browser, where different colors will distinguish T1- and T2- LADs. Alternatively, the user can separate T1- from T2-LADs using the values in column 4.

**Table 3: HMM-identified KDDs in the LAD atlas.** T1- and T2-KDDs for each cell type in the atlas in .bed format. Cell types separated by worksheets. All coordinates are in hg38. Each sheet can be exported into a .bed file and uploaded to a genome browser, where different colors will distinguish T1- and T2- KDDs. Alternatively, the user can separate T1- from T2-KDDs using the values in column 4.

**Table 4: GO Categories for Invariant T1- and T2-LADs.** Enriched gene ontology terms for genes located in invariants T1- or T2-LADs using the HumanBase Modules tool.

**Table 5: FISH probes locations.** Chromosomal regions used to design FISH probes provided. All probes were designed as tiled 80-mers with a density of 4probes/kb. Coordinates provided align to the hg38 genome assembly. Probes were amplified with direct dye conjugation (Cy2, Cy3, or Cy5), with the exception of the nonLAD probe, for which a fluorescent secondary probe was used (see Methods).

**Table 6: qPCR primer sequences used for LB1 and H3K9me2 HMM validation.** Sequences provided for the 20 T1-, T2- and nonLAD regions assessed for LB1 enrichment by ChIP-qPCR (Supp. Fig. 3C, see Methods) and the 20 T1- and T2-KDD regions assessed for H3K9me2 enrichment by ChIP-qPCR (Supp. Fig. 5D). Note: nonLAD primers were also used as nonKDD primers in Supp. Fig. 5D.

