## Supplementary figures and images for "An atlas of lamina-associated chromatin across twelve human cell types reveals an intermediate chromatin subtype"

### KeoughandShah_SuppFigs

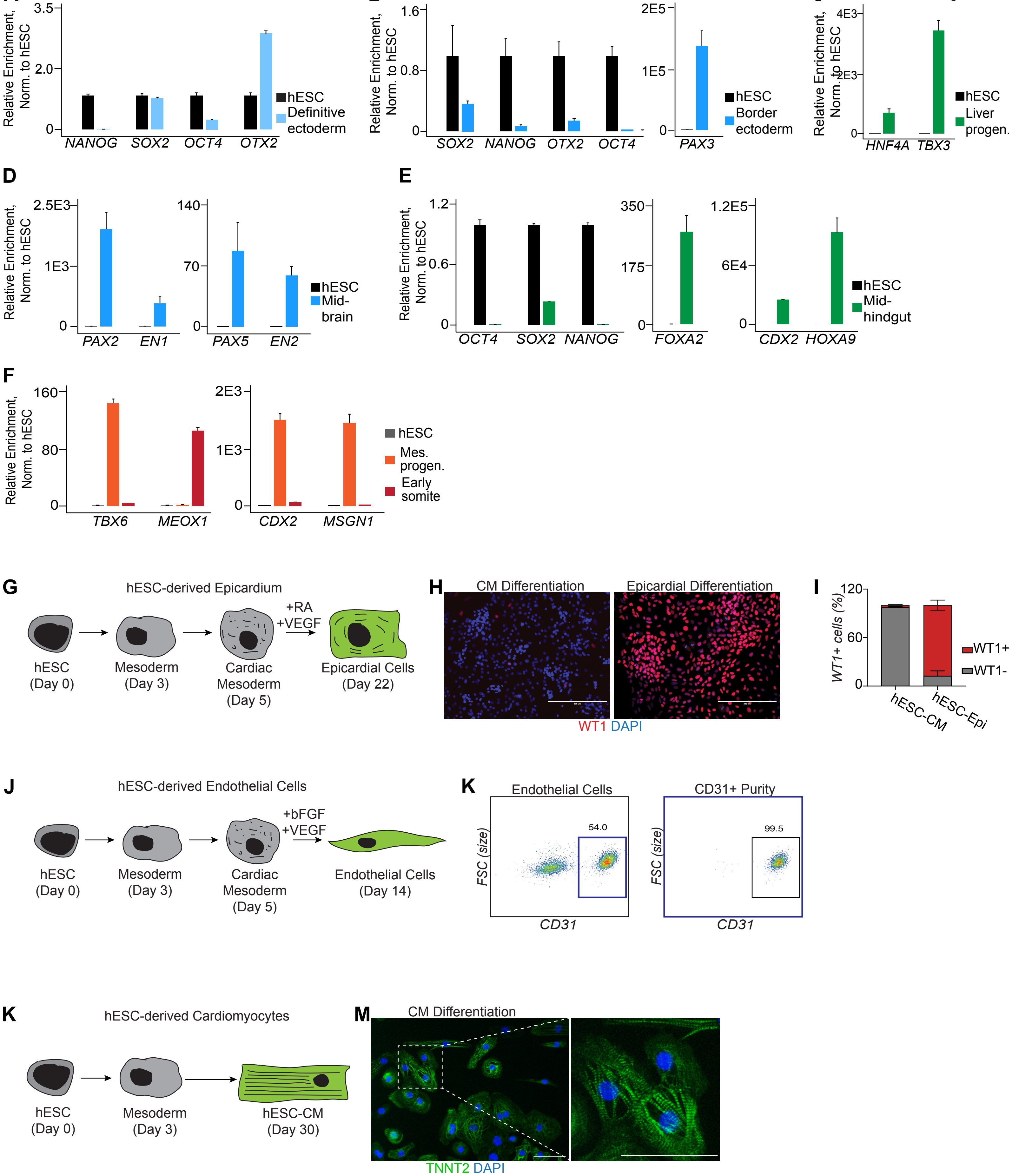

**S.Figure 2**

**A**

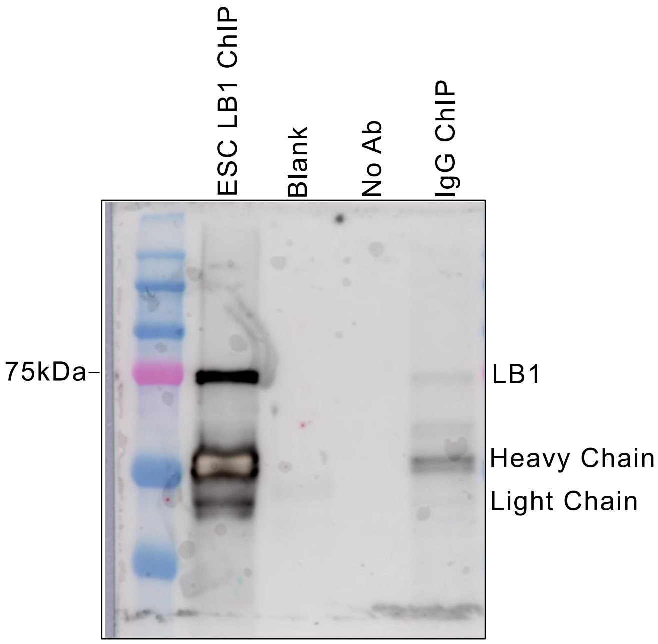


ChIP and IB: ab16048

**B**

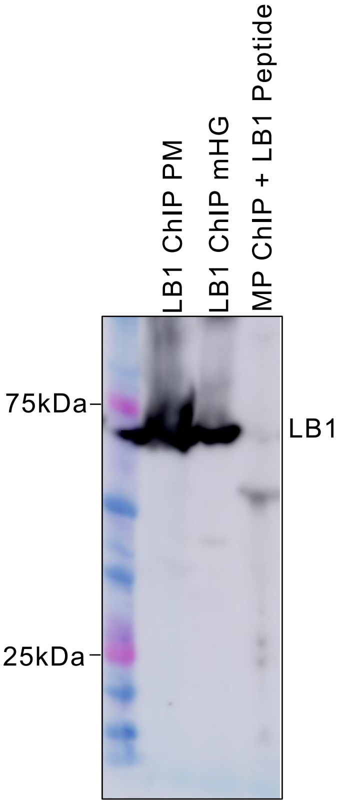


ChIP: ab16048
IB: sc-374015

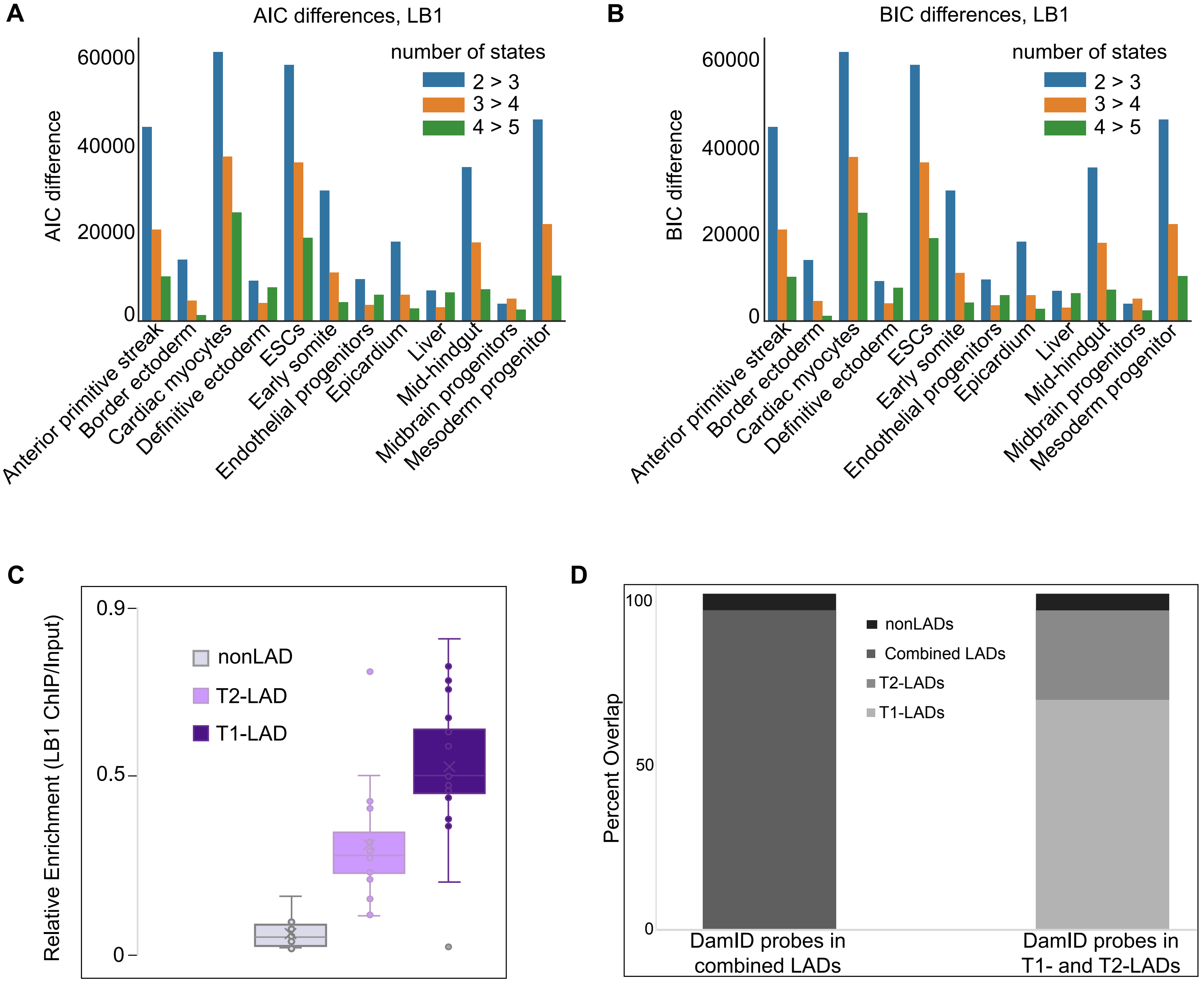

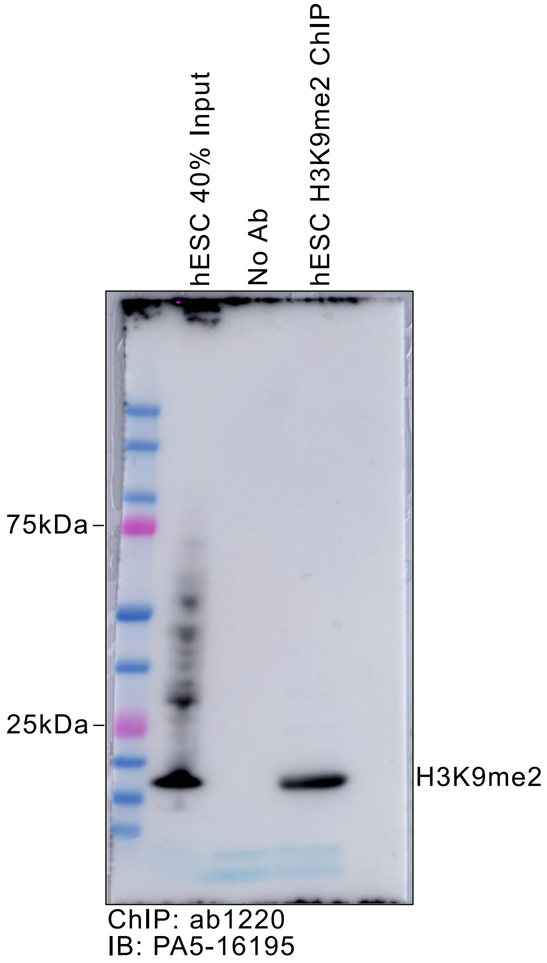

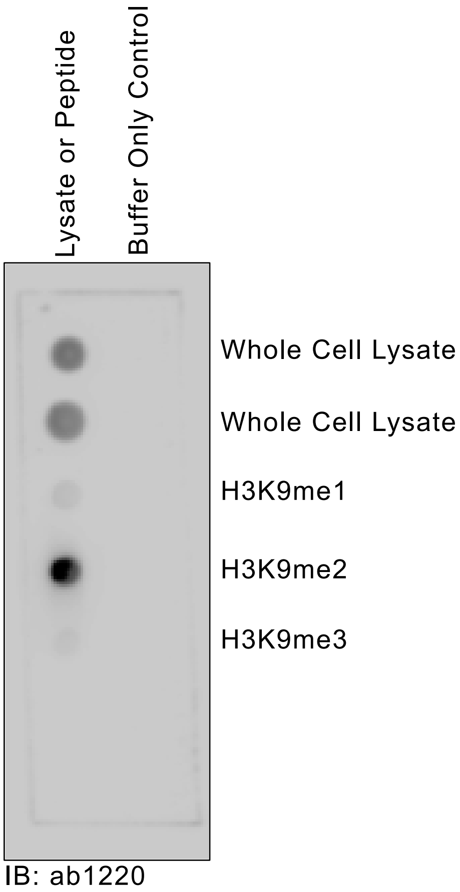

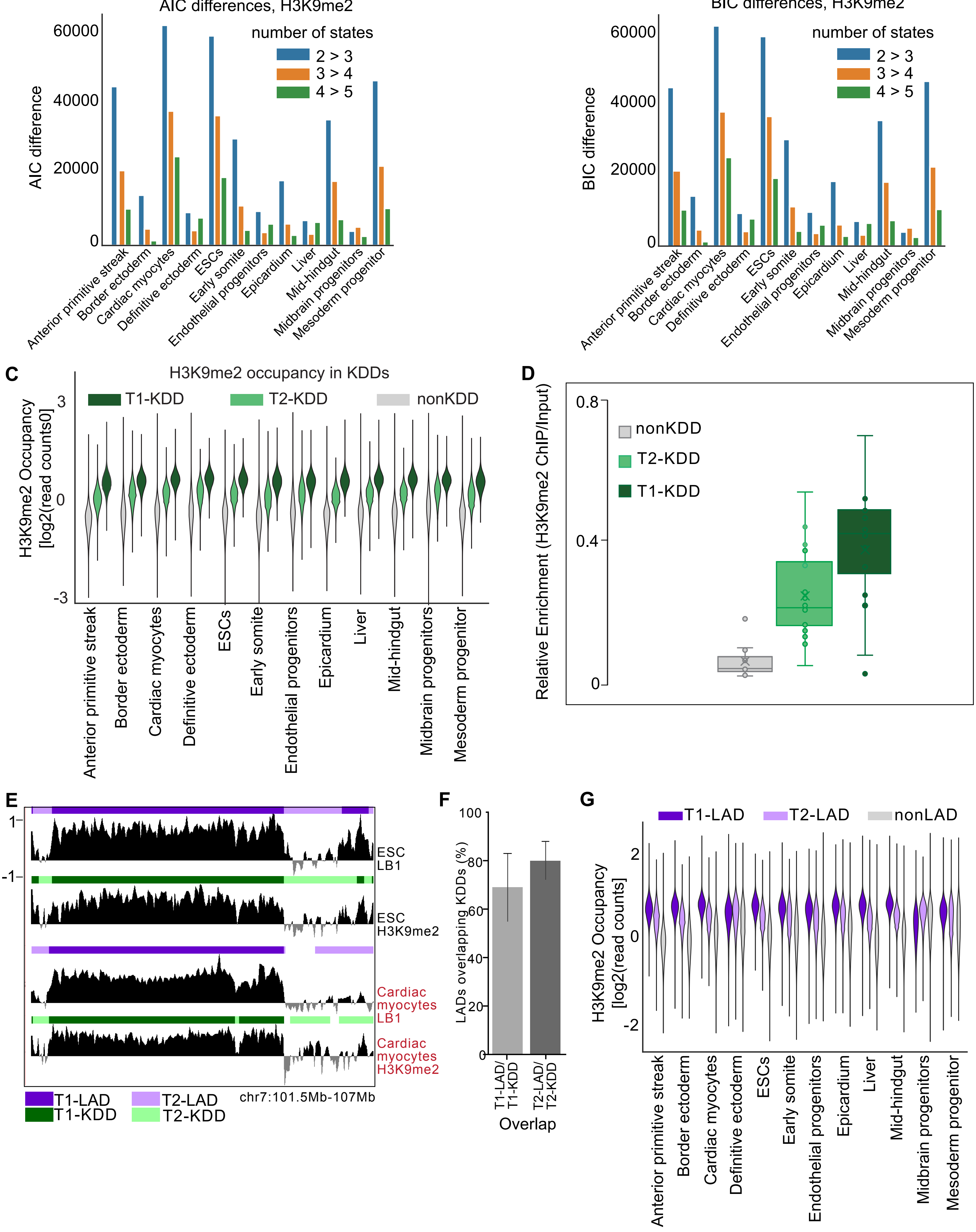

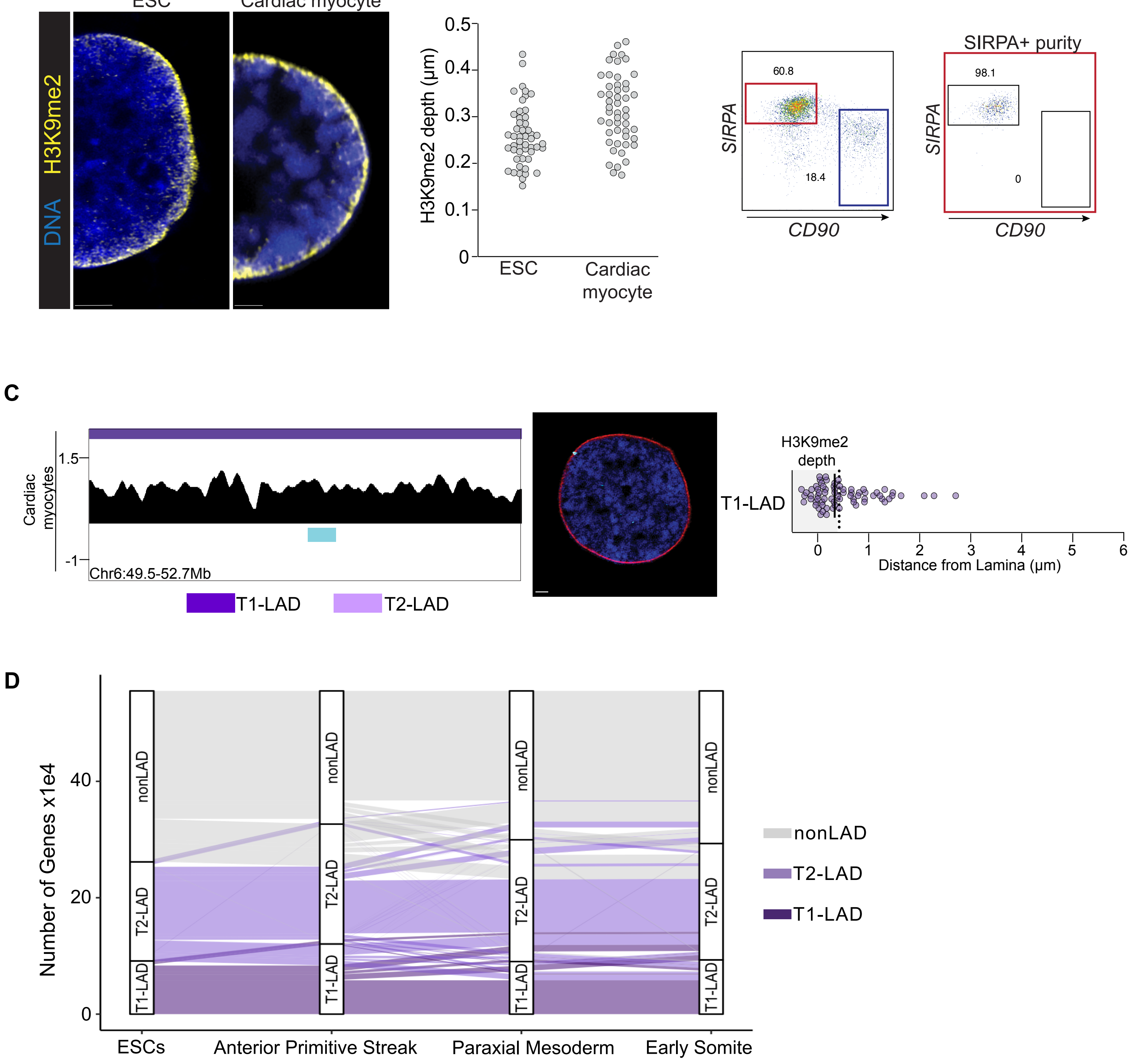
